# Genetic Encoding of Three Distinct Noncanonical Amino Acids Using Reprogrammed Initiator and Nonsense Codons

**DOI:** 10.1101/2020.12.07.415521

**Authors:** Jeffery M. Tharp, Oscar Vargas-Rodriguez, Alanna Schepartz, Dieter Söll

## Abstract

We recently described an orthogonal initiator tRNA (*i*tRNA^Ty2^) that can initiate protein synthesis with noncanonical amino acids (ncAAs) in response to the UAG nonsense codon. Here we report that a mutant of *i*tRNA^Ty2^ (*i*tRNA^Ty2^_AUA_) can efficiently initiate translation in response to the UAU tyrosine codon, giving rise to proteins with an ncAA at their N-terminus. We show that, in cells expressing *i*tRNA^Ty2^_AUA_, UAU can function as a dual-use codon that selectively encodes ncAAs at the initiating position and tyrosine at elongating positions. Using *i*tRNA^Ty2^_AUA_, in conjunction with its cognate tyrosyl-tRNA synthetase and two mutually orthogonal pyrrolysyl-tRNA synthetases, we demonstrate that UAU can be reassigned along with UAG or UAA to encode two distinct ncAAs in the same protein. Furthermore, by engineering the substrate specificity of one of the pyrrolysyl-tRNA synthetases, we developed a triply orthogonal system that enables simultaneous reassignment of UAU, UAG, and UAA to produce proteins containing three distinct ncAAs at precisely defined sites. To showcase the utility of this system, we produced proteins containing two or three ncAAs, with unique bioorthogonal functional groups, and demonstrate that these proteins can be separately modified with multiple fluorescent probes.

**TOC Image:** 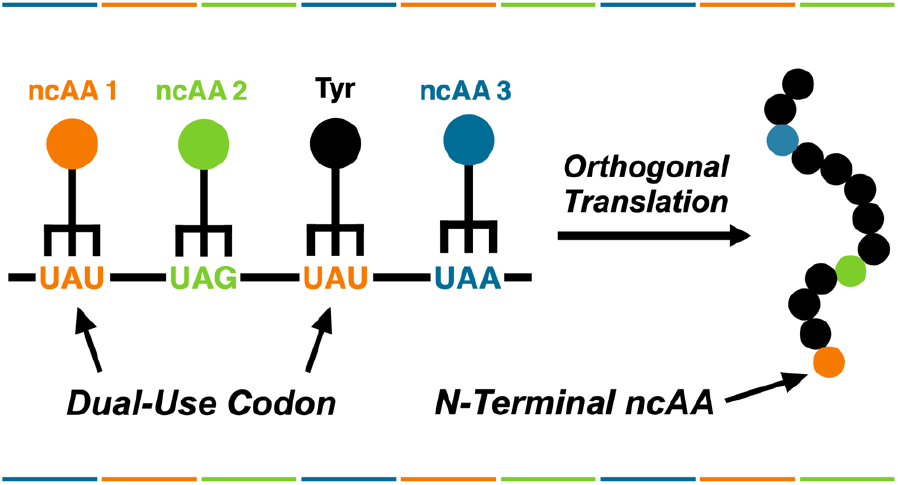

## INTRODUCTION

Most lifeforms on Earth utilize a ‘universal’ genetic code in which 64 triplet codons encode the twenty canonical amino acids and three stop signals. In the laboratory, however, artificial genetic code expansion (GCE) has enabled the biosynthesis of proteins containing diverse noncanonical amino acids (ncAAs) at precisely defined sites. This methodology employs orthogonal aminoacyl-tRNA synthetase (o-aaRS) and tRNA (o-tRNA) pairs to co-translationally install ncAAs into proteins, typically in response to a redefined nonsense codon. Laboratory GCE is not limited to the incorporation of a single ncAA. Through the combined action of multiple, mutually orthogonal o-aaRS•o-tRNA pairs, proteins with multiple distinct ncAAs can be produced in living cells.^1–5^ This technology has broad-ranging applications, from introducing reactive moieties into proteins for site-specific bioconjugation at multiple sites^4, 6^, to producing homogenously modified proteins featuring genetically encoded posttranslational modifications^7, 8^. An exciting prospect of GCE is the ability one day to produce completely unnatural polypeptides with new-to-Nature functions. However, further expansion of the genetic code is limited by the lack of ‘blank’ codons to encode additional unnatural moieties.

Most commonly, GCE relies on stop codon suppression, wherein a nonsense codon (i.e., UAG, UAA, or UGA) is reassigned to encode an ncAA. Recently, mutually orthogonal o-aaRS•o-tRNA pairs were used to simultaneously reassign all three nonsense codons and direct site-specific installation of three distinct ncAAs for the first time *in vivo*.^4^ However, nonsense suppression can only provide three blank codons for GCE. Moreover, reassigning all three nonsense codons requires modified expression systems that assist in accurately terminating translation.^4^ Four-base codons such as UAGA or AGGA represent a promising alternative with the possibility of providing 256 unique blank codons.^9^ Indeed, four-base codons have been used alone, and in combination with nonsense codons, to encode two and three distinct ncAAs in the same protein gene.^1, 5, 10^ However, translation of four-base codons is very inefficient on wildtype ribosomes.^3, 11, 12^ It is likely for this reason that four-base codons have not been widely adopted. A third strategy to further expand the genetic code is to reassign one of the 61 sense codons that normally encode a canonical amino acid.^13, 14^ This strategy is particularly challenging due to competition with endogenous aminoacyl-tRNAs for suppression of sense codons.^15^ Because of this competition, sense codon reassignment most often affords a heterogeneous product containing a mixture of the canonical and noncanonical amino acids.^14, 16–24^

In a recent study, we reported the development of an orthogonal initiator tRNA (*i*tRNA^Ty2^) that is a substrate for the *Methanocaldococcus jannaschii* tyrosyl-tRNA synthetase (*Mj*TyRS)—an o-aaRS that is commonly used for GCE.^25^ We demonstrated that *i*tRNA^Ty2^ can initiate translation at UAG codons with a variety ncAAs. The unique ability of *i*tRNA^Ty2^ to initiate translation is afforded by a conserved set of sequence motifs that are exclusive to initiator tRNAs.^25, 26^ Here, we asked whether *i*tRNA^Ty2^ could be engineered to initiate translation at a reassigned sense codon. We hypothesized that endogenous elongator tRNAs, which lack structural motifs required for initiation, would be unable to compete with *i*tRNA^Ty2^ for suppression of a reassigned initiator codon, thus abrogating a major hinderance to sense codon reassignment. We demonstrate that an anticodon mutant of *i*tRNA^Ty2^ can efficiently initiate translation with ncAAs in response to the UAU tyrosine codon. Initiation at UAU was achieved without detectable tyrosine incorporation giving rise to proteins with an ncAA at the N-terminus. Using this mutant initiator tRNA, alongside two mutually orthogonal pyrrolysyl-tRNA synthetase (PylRS) and tRNA pairs, we demonstrate that UAU can be reassigned along with UAA and UAG to simultaneously encode two and three distinct ncAAs in the same protein. Finally, we demonstrate that proteins containing two and three reactive ncAAs can be separately modified with multiple fluorescent probes.

## RESULTS AND DISCUSSION

### Reassigning sense codons for translation initiation with noncanonical amino acids

To investigate whether anticodon mutants of *i*tRNA^Ty2^ can initiate translation at reassigned sense codons, we employed a fluorescence-based assay that we previously used for measuring initiation at UAG.^25^ This assay relies on a superfolder green fluorescent protein (sfGFP)^27^ reporter in which the initiating AUG codon is replaced with UAG (sfGFP[1UAG]). The reporter is co-expressed with *i*tRNA^Ty2^ and a polyspecific *Mj*TyrRS variant (pCNFRS^28^ or AzFRS.2.t1^29^). Under these conditions, sfGFP production relies on the ability of *Mj*TyrRS to charge *i*tRNA^Ty2^ with an ncAA (provided in the growth media, **Figure 1**), and on the ability of aminoacyl-*i*tRNA^Ty2^ to initiate translation at UAG. To modify this assay for measuring initiation at sense codons, we constructed a series of reporter plasmids in which the initiating codon of sfGFP was replaced with one of eight sense codons (**Figure 2A**). Because *Mj*TyrRS interacts with the anticodon of its cognate tRNA, we limited our search to codons that required minimal base substitutions in *i*tRNA^Ty2^.^30^ First, to assess whether endogenous *E. coli* tRNAs can compete for initiation at these codons, we measured the expression of each sfGFP reporter in the absence of an *i*tRNA^Ty2^ mutant. Without co-expression of a mutant initiator tRNA very little sfGFP expression was detected with each reporter (**Figure 2B**), supporting our assumption that endogenous *E. coli* tRNAs cannot initiate translation at these sense codons.

**Figure 1.**
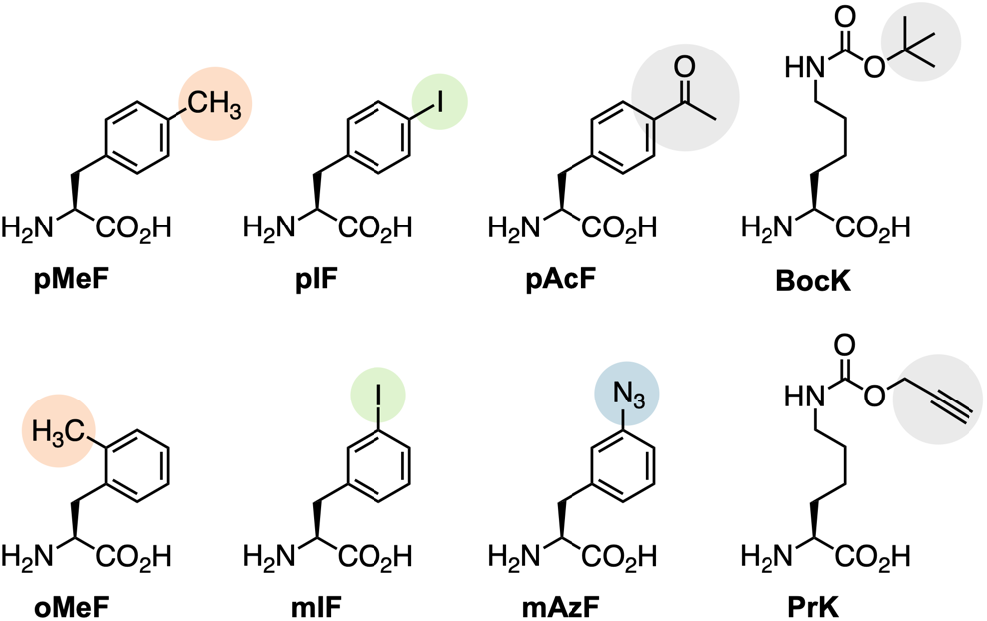
The structures of ncAAs used in this study.

**Figure 2.**
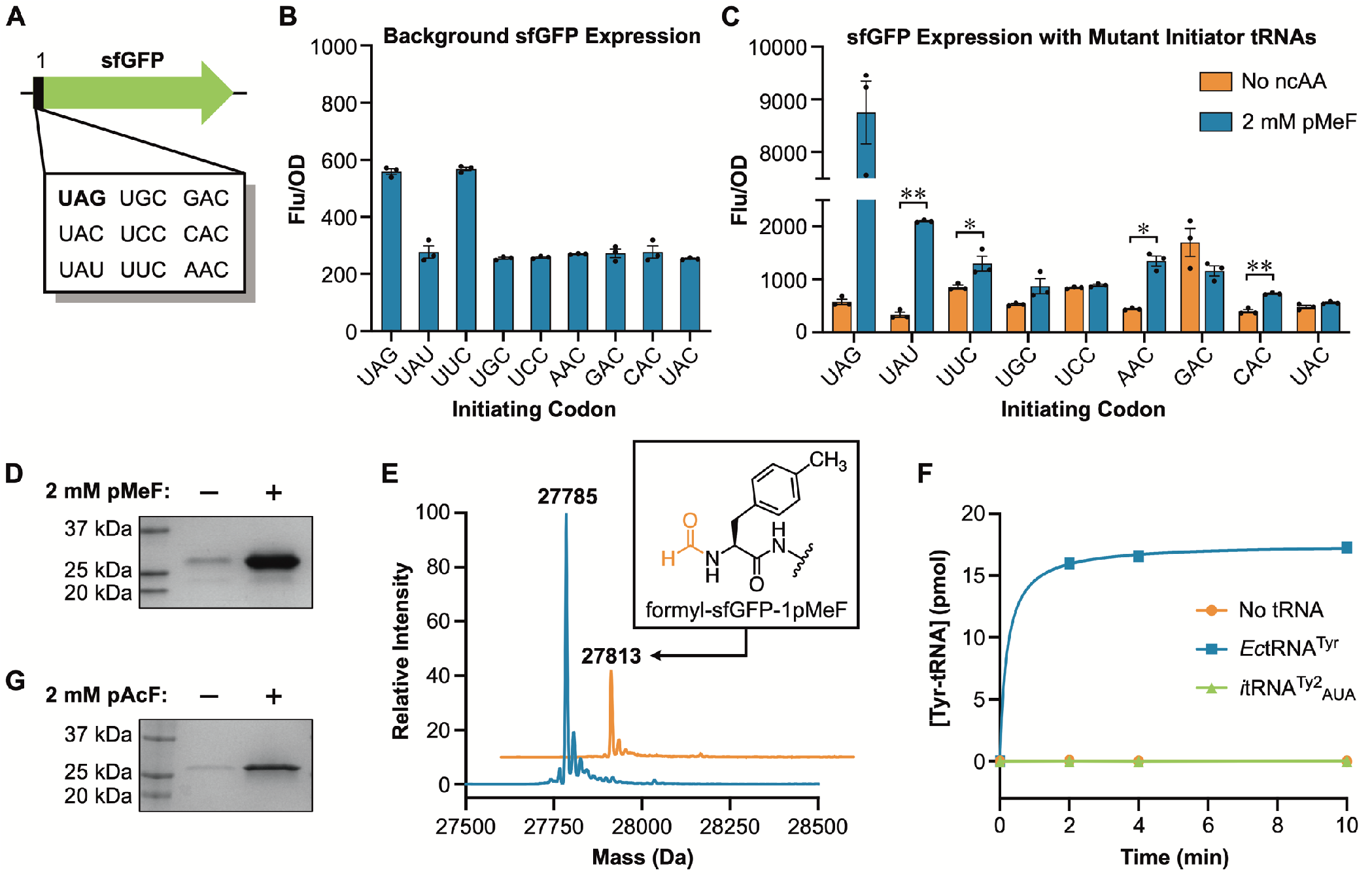
Reassigning sense codons for initiation with ncAAs. (A) A series of sfGFP reporters were constructed in which the initiating methionine codon was replaced with UAG or one of eight sense codons. (B) Endogenous *E. coli* tRNAs cannot initiate at UAG or any of the sense codons. Data are displayed as the mean ± SEM of three biological replicates. (C) Co-expression of a mutant initiator tRNA enables translation initiation with ncAAs at select codons. Data are displayed as the mean ± SEM of three biological replicates (* = *p* < 0.05, ** = *p* < 0.005, paired t-test). (D) Expression of sfGFP[1UAU] is dependent on addition of pMeF to the growth media. (E) LC-MS of D. Peaks corresponding to sfGFP-1pMeF (theoretical mass = 27786 Da) and *N*-formyl-sfGFP-1pMeF (theoretical mass = 27814 Da) were detected. (F) *i*tRNA^Ty2^_AUA_ is not aminoacylated by the *E. coli* TyrRS. Timepoints represent the mean of three independent experiments. (G) ncAA-dependent expression of codon-optimized sfGFP in which all elongating UAU codons were replaced with UAC.

Next, we measured the expression of each sfGFP mutant in *E. coli* cells that were simultaneously expressing *i*tRNA^Ty2^ (with complementary anticodon mutations) and pCNFRS. With four of the initiating sense codons (UAU, UUC, AAC, CAC) we observed significant sfGFP expression when the ncAA *para*-methyl-L-phenylalanine (pMeF) was added to the growth media (**Figures 1** and **2C**). This indicates that the *i*tRNA^Ty2^ mutants were successfully charged with pMeF and that the aminoacyl-tRNA can initiate translation at the reassigned sense codon. Of the four codons that showed significant sfGFP expression with pMeF, the tyrosine codon UAU was most efficient, affording a fluorescence/OD_600_ signal ~24% as intense as UAG initiation (**Figure 2C**). We measured sfGFP[1UAU] expression in wildtype *E. coli* DH10B, and in an engineered strain in which redundant copies of the methionine initiator tRNA gene were deleted from the genome (strain DH10BΔ*metZWV*). We have shown that initiation with ncAAs is far more efficient in DH10BΔ*metZWV* than in wildtype cells.^25^ Consistent with our assumption that the mutant initiator tRNA (*i*tRNA^Ty2^_AUA_) is initiating translation at UAU, sfGFP[1UAU] expression was nearly five-fold greater in DH10BΔ*metZWV* than in the wildtype strain (**Figure S1**).

To confirm pMeF incorporation in response to UAU, we expressed and purified sfGFP[1UAU] using pCNFRS and *i*tRNA^Ty2^_AUA_. Consistent with in-cell fluorescence measurements, robust expression of the reporter only occurred in the presence of pMeF (**Figure 2D**and **S2A**). LC-MS of the purified protein revealed a major peak matching the expected mass of sfGFP with pMeF in place of the initiating methionine (sfGFP-1pMeF, **Figure 2E**). We also observed a peak corresponding to *N*-formyl-sfGFP-1pMeF (**Figure 2E**). We have shown that when ncAAs initiate translation in DH10B-Δ*metZWV* a significant fraction of sfGFP retains an N-terminal formyl modification.^25^ This result further supports the conclusion that pMeF-charged *i*tRNA^Ty2^_AUA_ is initiating translation at UAU. Importantly, no peaks corresponding to tyrosine incorporation at the initiating UAU codon were detected, further demonstrating that the endogenous *E. coli* tyrosine tRNA (*Ec*tRNA^Tyr^) is unable to initiate translation at UAU.

In a previous study we showed that *i*tRNA^Ty2^ can both initiate and elongate translation at UAG codons.^25^ In addition to the initiating UAU, our sfGFP reporter has five elongating UAU codons; however, pMeF was not detected at these positions. This could be due to the low mass difference between pMeF and tyrosine (Δmass = 1.97 Da) which makes it difficult to distinguish between these residues based on mass alone. Therefore, we expressed sfGFP[1UAU] with *para*-iodo-L-phenylalanine (pIF; **Figure 1**) which has a larger mass than tyrosine (Δmass = 109.9 Da). Intact LC-MS of the purified protein revealed major peaks corresponding to sfGFP-1pIF and *N*-formyl-sfGFP-1pIF (**Figure S3**). No peaks corresponding to tyrosine incorporation at the initiating UAU were detected. Likewise, no peaks corresponding to pIF incorporation at elongating UAU codons were detected. We further analyzed this protein by tandem mass spectrometry (MS/MS) following proteolysis. In this analysis we identified peptides with masses and fragmentation patterns consistent with pIF incorporation at the N-terminal position (**Figure S4**); we were unable to detect pIF incorporation in response to any elongating UAU codon. Thus, our data suggest that while *i*tRNA^Ty2^_AUA_ can initiate translation with ncAAs at UAU, under these conditions, it is unable to compete with endogenous *Ec*tRNA^Tyr^ for suppression of elongating UAU codons. Taken together, our results indicate that, in cells expressing *i*tRNA^Ty2^_AUA_, UAU can function as a dual-use codon: at the initiating position UAU encodes an ncAA, whereas at elongating positions UAU encodes tyrosine. Similar dual-use codons have been reported *in vitro*^31^ and partial dual reassignment of the methionine AUG codon has been reported *in vivo*^18^; however, to our knowledge, this is the first time that complete reassignment of a dual-use codon has been achieved in living cells. It should be noted that this observation might be codon context dependent. That is, incorporation of ncAAs at elongating UAU codons cannot be ruled out for every protein at this time. To avoid possible complications from elongating UAU codons in subsequent experiments, we synthesized a codon-optimized sfGFP in which all elongating UAU codons were replaced with the synonymous codon UAC. As with our previous reporter, robust expression of the codon-optimized variant occurred only when an ncAA was provided in the growth media (**Figure 2G** and **S2B**). When this optimized gene was expressed with *para*-acetyl-L-phenylalanine (pAcF; **Figure 1**), sfGFP-1pAcF was obtained with a yield of ~260 mg per liter of culture.

Several of the codons that we tested showed an increase in sfGFP expression irrespective of whether an ncAA was provided in the growth media (**Figure 2C**). A possible explanation for this observation is that mutating the anticodon of *i*tRNA^Ty2^ converted the tRNA into a substrate for an endogenous *E. coli* aaRS. For example, we observed an increase in sfGFP expression when the anticodon of *i*tRNA^Ty2^ was mutated to GUC (for initiating at GAC). In *E. coli* GAC encodes aspartate and the aspartyl-tRNA synthetase uses anticodon bases to recognize its cognate tRNA.^32^ Similar misaminoacylation was observed when the anticodon of pyrrolysine tRNA (tRNA^Pyl^) was mutated to reassign an arginine codon in *Mycoplasma capricolum*.^16^ While no tyrosine incorporation at the initiating UAU codon was detected, it is possible that *i*tRNA^Ty2^_AUA_ is still being mischarged with tyrosine by *E. coli* TyrRS (*Ec*TyrRS). Proteins that contain an N-terminal tyrosine are extremely unstable in *E. coli*^33^, thus, mis-charging of *i*tRNA^Ty2^_AUA_ with tyrosine might go undetected if the reporter protein is rapidly degraded *in vivo*. To investigate this possibility, we performed *in vitro* aminoacylation assays using purified *Ec*TyrRS and an *i*tRNA^Ty2^_AUA_ transcript. While *Ec-*TyrRS charged its cognate tRNA with radiolabeled tyrosine, no aminoacylation of *i*tRNA^Ty2^_AUA_ was detected (**Figure 2F**), indicating that the mutant tRNA is orthogonal to *Ec*TyrRS.

### Dual incorporation of noncanonical amino acids using reprogrammed initiator and nonsense codons

After demonstrating that *i*tRNA^Ty2^_AUA_ can initiate translation with ncAAs at UAU codons, we next asked whether UAU could be used together with nonsense codons to encode two distinct ncAAs in the same protein. The two most commonly used codons for GCE are UAG and UAA; however, *i*tRNA^Ty2^_AUA_ can base pair with these triplets at all but the wobble position. To encode distinct ncAAs using a combination of UAU and UAG/UAA, *i*tRNA^Ty2^_AUA_ must be able to recognize UAU and discriminate against UAG and UAA. Therefore, to evaluate whether *i*tRNA^Ty2^_AUA_ recognizes these nonsense codons, we constructed two reporters in which the second position of sfGFP was mutated to UAG or UAA (sfGFP[2UAG] and sfGFP[2UAA], respectively; **Figure S5A**). We co-expressed these mutant reporters, along with *i*tRNA^Ty2^_AUA_ and AzFRS.2.t1, in *E. coli* DH10BΔ*metZWV*, and we monitored sfGFP production in the presence of pMeF. As expected, sfGFP[1UAU] afforded a strong fluorescence signal, whereas no significant fluorescence was observed in cells expressing sfGFP[2UAG] or sfGFP[2UAA] (**Figure S5B**), demonstrating that *i*tRNA^Ty2^_AUA_ is orthogonal to these nonsense codons.

After confirming orthogonality of *i*tRNA^Ty2^_AUA_ towards UAG and UAA, we next explored whether these codons could be used, together with a UAU, to encode two different ncAAs. For introducing a second ncAA we conscripted the PylRS•tRNA^Pyl^ pair which is orthogonal in bacteria and eukaryotes and has been engineered to recognize numerous ncAA substrates.^34, 35^ The most widely used PylRS•tRNA^Pyl^ pairs for GCE are those originating from the *Methanosarcina* species *mazei* (*Mm*PylRS•*Mm*tRNA^Pyl^) and *barkeri*. Importantly, the PylRS and *Mj*TyrRS systems are mutually orthogonal and have been used to install distinct ncAAs in response to UAA and UAG codons within the same gene.^2^ Here, we asked whether the *Mj*TyrRS and *Mm*PylRS can be used to introduce two unique ncAAs in response to UAU and UAA, respectively. With this objective we assembled a plasmid system for simultaneous expression of the two o-aaRS•o-tRNA pairs, together with a sfGFP reporter containing a UAU mutation at position 1 and a UAA mutation at position 151 (sfGFP[1UAU-151UAA]; **Figure S6A**). In this system we employed the *Mj*TyrRS variant AzFRS.2.t1, together with *i*tRNA^Ty2^_AUA_, to incorporate pMeF in response to the initiating UAU and wildtype *Mm*PylRS, together with a mutant opal-suppressor *Mm*tRNA^Pyl^ (*Mm*tRNA^Pyl^_UUA_)^36^, to introduce *N*^ε^-boc-L-lysine (BocK; **Figure 1**) in response to the elongating UAA. We measured sfGFP production in *E. coli* DH10BΔ*metZWV* in the presence of both pMeF and BocK. Under these conditions, robust expression occurred only when both ncAAs were provided in the growth media (**Figure 3A**), suggesting that the UAU and UAA codons were simultaneously suppressed to afford full-length sfGFP with the desired ncAAs.

**Figure 3.**
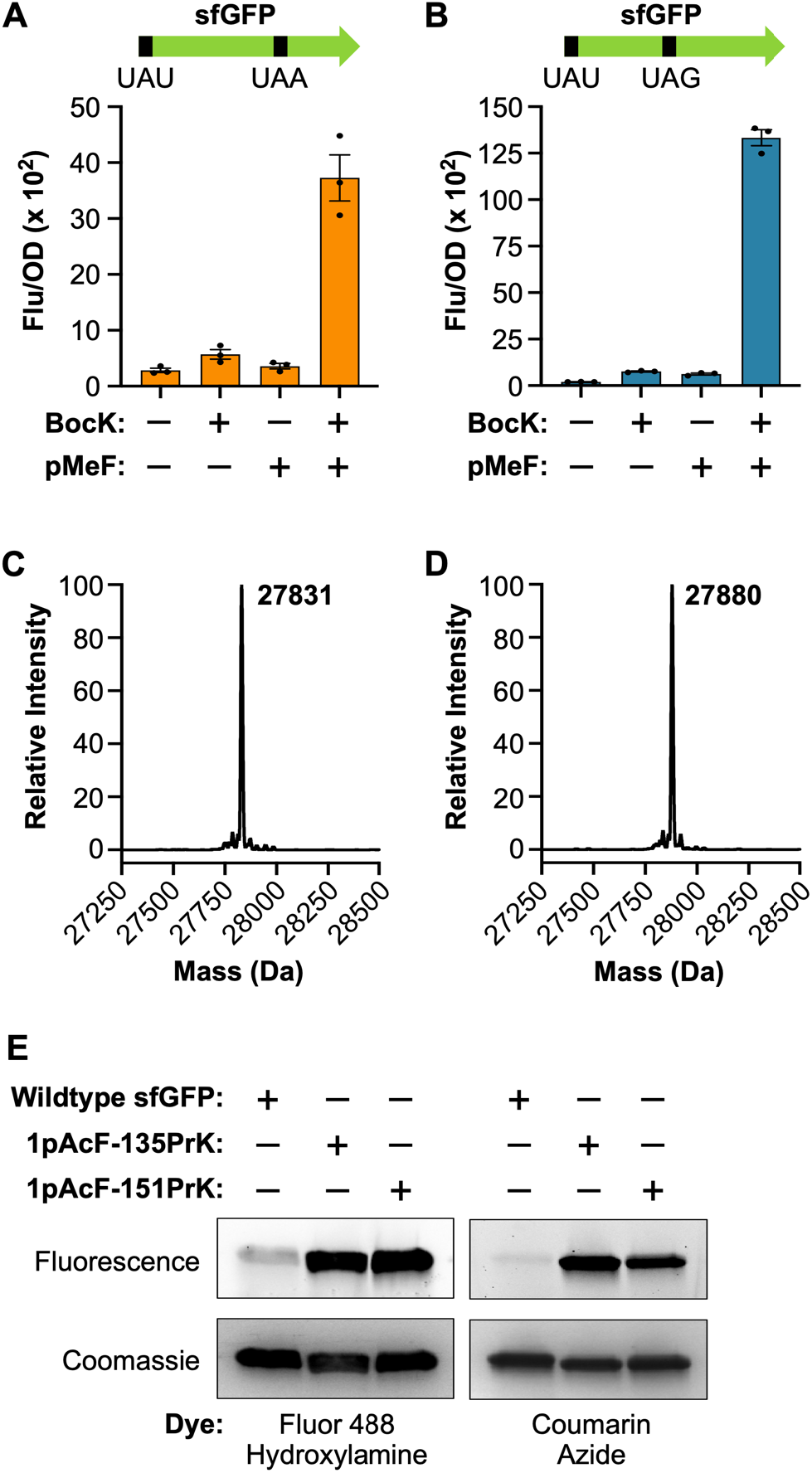
Co-translational installation of two distinct ncAAs encoded by UAU/UAA and UAU/UAG. (A-B) Expression of sfGFP[1UAU-151UAA] (A) and sfGFP[1UAU-135UAG] (B) in the presence of pMeF and BocK. Data are displayed as the mean ± SEM of three biological replicates. (C) LC-MS of sfGFP-1pAcF-135PrK (theoretical mass = 27833 Da). (D) LC-MS of sfGFP-1pAcF-151PrK (theoretical mass = 27882 Da). (E) Labeling of sfGFP-1pAcF-135PrK and sfGFP-1pAcF-151PrK with Fluor 488-hydroxylamine and coumarin azide. Proteins were resolved by SDS-PAGE and visualized by ingel fluorescence and Coomassie staining.

Next, we evaluated dual ncAA incorporation using a com-bination of UAU and UAG. To introduce a second ncAA in response to UAG, we employed a recently characterized PylRS•tRNA^Pyl^ pair from the methanogenic archaeon “*Candi-datus* Methanomethylophilus alvus” (*Ma*PylRS•*Ma-*tRNA^Pyl^).^37^ Like the *Mm*PylRS pair, the *Ma*PylRS•*Ma*tRNA^Pyl^ pair is orthogonal in bacteria and eukaryotes.^37, 38^ Furthermore, we have shown that this pair is mutually orthogonal with the engineered *Mj*TyrRS•*i*tRNA^Ty2^ pair.^25^ We employed the same plasmid-based system to co-express these two o-aaRS•o-tRNA pairs along with a sfGFP mutant containing a UAU mutation at position 1 and a UAG mutation at position 135 (sfGFP-[1UAU-135UAG]; **Figure S6B**). AzFRS.2.t1 and *i*tRNA^Ty2^_AUA_ were used to incorporate pMeF in response to the initiating UAU, while wildtype *Ma*PylRS and a variant of *Ma*tRNA^Pyl^ (*Ma*tRNA(6)^Pyl^), which was engineered to be orthogonal to *Mm*PylRS, were used to incorporate BocK in response to UAG.^37^ Again, robust sfGFP expression occurred only when both pMeF and BocK were added to the growth media, demonstrating dual suppression of UAU and UAG and successful incorporation of both ncAAs (**Figure 3B**).

AzFRS.2.t1 and the wildtype PylRSs are polyspecific synthetases that recognize a number of substrates.^29, 34^ Among these are ncAAs with bioorthogonal functional groups that enable site-specific protein bioconjugation. For example, in addition to pMeF, AzFRS.2.t1 recognizes the ketone-containing ncAA pAcF which can undergo oxime ligation with hydroxylamine probes.^39^ Furthermore, in addition to BocK, wildtype PylRS recognizes *N*^ε^-propargyl-L-lysine (PrK; **Figure 1**) which can undergo Cu(I)-catalyzed cycloaddition with azides.^40^ We tested whether these o-aaRS•o-tRNA pairs could be used to produce proteins containing ketone and alkyne reactive moieties at precisely defined sites by expressing sfGFP[1UAU-135UAG] and sfGFP[1UAU-151UAA] in the presence of pAcF and PrK. Again, robust expression of these reporter proteins occurred only when both ncAAs were provided in the growth media (**Figure S6C–D**). Next, we purified these proteins and confirmed pAcF and PrK incorporation by LC-MS; in both cases we observed mass peaks consistent with sfGFP containing the expected ncAA substitutions (**Figure 3C–D**). We also confirmed the site-specificity of pAcF and PrK incorporation by MS/MS analysis (**Figure S7–8**). Finally, to demonstrate the utility of this system, we showed that these proteins, which contain ketone and alkyne bioconjugation handles, can be readily labeled with both hydroxylamine- and azide-based fluorescent dyes (**Figure 3E** and **S9**).

### Co-translational installation of three distinct noncanonical amino acids

The above data demonstrate that *Mj*TyrRS•*i*tRNA^Ty2^_AUA_ can be used, in conjunction with two different PylRS•tRNA^Pyl^ pairs, to incorporate distinct ncAAs in response to UAU and UAG/UAA. Next, we asked whether these mutually orthogonal o-aaRS•o-tRNA pairs can be combined to direct site-specific installation of three unique ncAAs. As a prerequisite, we first sought to identify three o-aaRS variants that recognize distinct ncAAs substrates. The *Mj*TyrRS variant AzFRS.2.t1 primarily recognizes *para*-substituted phenylalanine derivatives, whereas wildtype *Mm*PylRS and *Ma*PylRS both recognize *N*^ε^-substituted lysine derivatives and have overlapping substrate specificities.^41^ A major contributor to the substrate specificity of PylRS is the so-called *gatekeeper residue*, a conserved asparagine located in the enzyme’s substrate binding pocket. This residue is involved in several interactions with the substrate amino acid, including a hydrogen bond with the side chain amide oxygen of pyrrolysine, and other *N*^ε^-substituted lysine derivatives (**Figure S10A**).^42, 43^ Mutating this asparagine in *Mm*PylRS to alanine (N346A) obliterates recognition of *N*^ε^-substituted lysine derivatives.^44^ Introducing a second alanine mutation (C348A) affords an *Mm*PylRS variant (*Mm*PylRS(N346A/C348A)) that recognizes more than thirty *ortho*-, *para*-, and *meta*-substituted phenylalanine derivatives.^44–46^ However, this variant poorly recognizes derivatives with small *para* substitutions, such as those recognized by AzFRS.2.t1.^44^ Therefore, we hypothesized that *Mm*PylRS(N346A/ C348A) and AzFRS.2.t1 could be used together in the same cell to selectively install *meta*- and *para*-substituted phenylalanine derivatives, respectively. In addition, we hypothesized that wildtype *Ma*PylRS could serve as a third o-aaRS to install *N*^ε^-substituted lysine derivatives.

To confirm the substrate specificity of these o-aaRSs, we measured sfGFP[2UAG] production in cells expressing either AzFRS.2.t1, *Mm*PylRS(N346A/C348A), or wildtype *Ma*PylRS (as well as their cognate amber-suppressor tRNAs) in the presence of three substrates: pAcF, PrK, and *meta*-iodo-L-phenylalanine (mIF; **Figure 1**). Consistent with our hypothesis, we found that AzFRS.2.t1 selectively recognizes the *para*-substituted ncAA pAcF, while *Mm*PylRS(N346A/C348A) selectively recognizes the *meta*-substituted ncAA mIF (**Figure 4A–B**). However, surprisingly, we found that mIF is also recognized by wildtype *Ma*PylRS (**Figure 4C**). Indeed, *Ma*PylRS enabled robust sfGFP expression in the presence of both mIF and PrK. In contrast, wildtype *Mm*PylRS only afforded robust reporter expression in the presence of PrK (**Figure 4B**).

**Figure 4.**
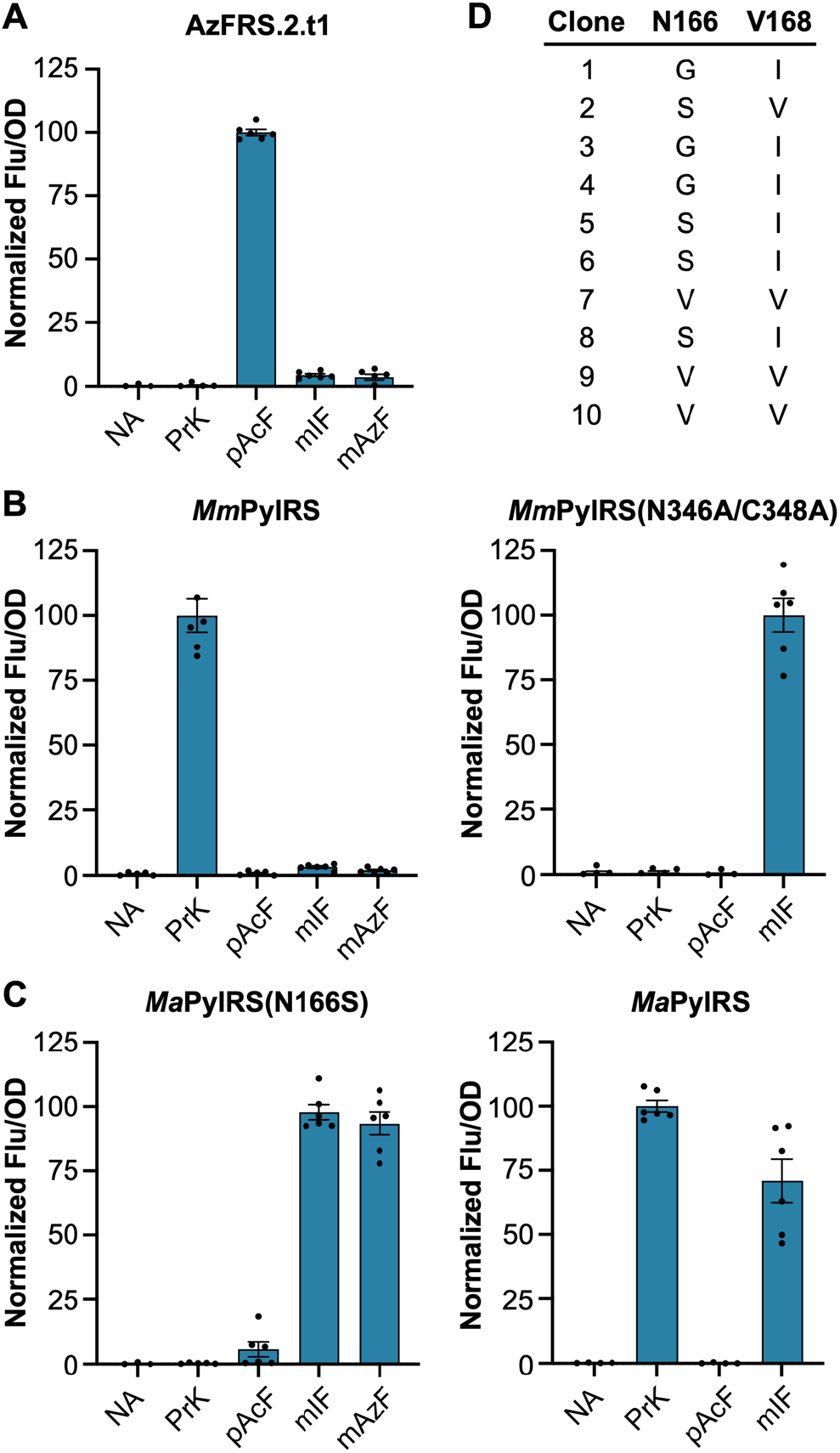
Substrate specificities of three mutually orthogonal aminoacyl-tRNA synthetases. (A-C) Expression of sfGFP[2UAG] in the presence of PrK, pAcF, mIF, or mAzF facilitated by (A) the *Mj*TyrRS variant AzFRS.2.t1, (B) wildtype *Mm*PylRS and variant *Mm*PylRS(N346A/C348A), and (C) wildtype *Ma*PylRS and variant *Ma*PylRS(N166S) . Data are displayed as the mean ± SEM of six biological replicates. NA = None Added. (D) *Ma*PylRS mu-tants identified from an N166/V168 library selected on media containing mIF (clones 1–5) or oMeF (clones 6–10) .

In light of this observation, we decided instead to use *Ma*PylRS to install *meta*-substituted ncAAs and wildtype *Mm*PylRS to install *N*^ε^-substituted lysine derivatives. However, this required that we engineer an *Ma*PylRS variant that accepts mIF and rejects PrK. Towards this end, we constructed a two-site *Ma*PylRS library randomizing residues N166 and V168 (corresponding to N346 and C348 in *Mm*PylRS, **Figure S10B**). We subjected this library to positive selection using chloramphenicol acetyltransferase with a UAG codon at position 112 (cat[112UAG]).^47^ Cells expressing the *Ma*PylRS N166/V168 library, *Ma*tRNA(6)^Pyl^, and cat[112UAG] were challenged to grow on media containing 1 mM mIF and 50 μg/mL chloramphenicol. We also performed a parallel selection using *ortho*-methyl-L-phenylalanine (oMeF; **Figure 1**). Following selection, surviving clones were screened for ncAA-dependent growth (**Figure S11**) and five clones from each selection were sequenced to reveal mutations in *Ma*PylRS that altered its specificity. In total, four unique mutants with similar amino acid substitutions were identified (**Figure 4D**). We screened these mutants for selective incorporation of mIF using sfGFP[2UAG]. Gratifyingly, we found that robust expression occurred only in the presence of mIF, indicating that these newly identified *Ma*PylRS mutants are selective for the *meta*-substituted ncAA (**Figure 4C** and **S12**). In subsequent experiments we employed the N166S mutant, *Ma*PylRS(N166S). In addition to mIF and oMeF, we screened *Ma*PylRS(N166S) for the ability to recognize various substituted phenylalanine derivatives. Similar to *Mm*PylRS-(N346A/C348A), *Ma*PylRS(N166S) recognizes a number of substrates (**Figure S13**) including those with bioorthogonal handles such as *meta*-azido-L-phenylalanine (mAzF; **Figure 1**). Like mIF, we found that mAzF is rejected by AzFRS.2.t1 and wildtype *Mm*PylRS (**Figure 4A–C**). To test if this *ortho/para/meta* substrate selectivity is a general feature of *Ma*PylRS(N166S) and AzFRS.2.t1 we compared sfGFP[2UAG] expression with these enzymes using a panel of phenylalanine derivatives containing the same substituent at different positions of the aromatic ring. In all cases we found that AzFRS.2.t1 selectively recognizes *para*-substituted deri-vates, while *Ma*PylRS(N166S) selectively recognizes derivatives with *ortho* or *meta* substitutions (**Figure S14**).

Next, we devised a plasmid-based system for simultaneous incorporation of three distinct ncAAs. This three-plasmid system consisted of: (1) an accessory plasmids encoding AzFRS.2.t1, *Ma*PylRS(N166S), and *Ma*tRNA(6)^Pyl^, (2) a second accessory plasmid encoding *Mm*PylRS and *Mm*tRNA^Pyl^_UUA_, and (3) a reporter plasmid encoding *i*tRNA^Ty2^_AUA_ and a triple-mutant sfGFP with a UAU mutation at position 1, a UAG mutation at position 135, and a UAA mutation at position 151 (**Figure S15A**). We measured sfGFP production *E. coli* DH10BΔ*metZWV* in the presence of pMeF, BocK, and mIF. With this initial system, we observed significant sfGFP production only in the presence of all three ncAAs, however, the signal was relatively low (**Figure S15B**). We hypothesized that this was due to poor suppression of the UAA codon by *Mm*tRNA^Pyl^_UUA_ since, *Mm*PylRS is the least active o-aaRS in this system. Therefore, an additional copy of *Mm*tRNA^Pyl^_UUA_ was added to the reporter plasmid resulting in a ~2-fold increase in the fluorescence signal (**Figure S15B**). Next, we purified sfGFP expressed with pAcF, PrK, and mIF and we analyzed the protein by LC-MS. This analysis revealed a major peak consistent with sfGFP containing three ncAAs at the desired positions, however, a second peak was also present (**Figure S16**). This second peak was consistent with misincorporation of PrK at the UAG codon. Further analysis by MS/MS confirmed a mixture of mIF and PrK at position 135 **(Figure S17–18**). We surmised that mis-incorporation of PrK likely results from an excess of *Mm*tRNA^Pyl^_UUA_, since ochre-suppressor tRNAs can suppress both UAA and UAG.^48^ Therefore, we constructed a third reporter plasmid in which the extra copy of *Mm*tRNA^Pyl^_UUA_ was replaced with *Ma*tRNA(6)^Pyl^ (**Figure S15A**). Using this new plasmid, we again found that sfGFP production occurred only when the growth media was supplemented with a *para*-substituted phenylalanine, a *meta*-substituted phenylalanine, and an *N*^ε^-substituted lysine derivative, i.e., pMeF, mIF, and BocK or pAcF, mIF, and PrK. (**Figure 5A–B**). LC-MS analysis of sfGFP expressed with pAcF, mIF, and PrK revealed a major peak consistent with the incorporation of all three amino acids; a peak corresponding to misincorporation of PrK was not detected (**Figure 5C**). Furthermore, MS/MS analysis confirmed the incorporation of each ncAA at the desired positions (**Figure S19**). We used this optimized expression system to site-specifically install three reactive ncAAs (pAcF, mAzF, and PrK) into sfGFP affording a protein with three unique bioorthogonal functional groups (ketone, azide, and alkyne; **Figure 5D**). Finally, we demonstrated that this single protein could be modified with multiple fluorescent probes bearing hydroxylamine, alkyne, or azide reactive handles (**Figure 5D** and **S20)**.

**Figure 5.**
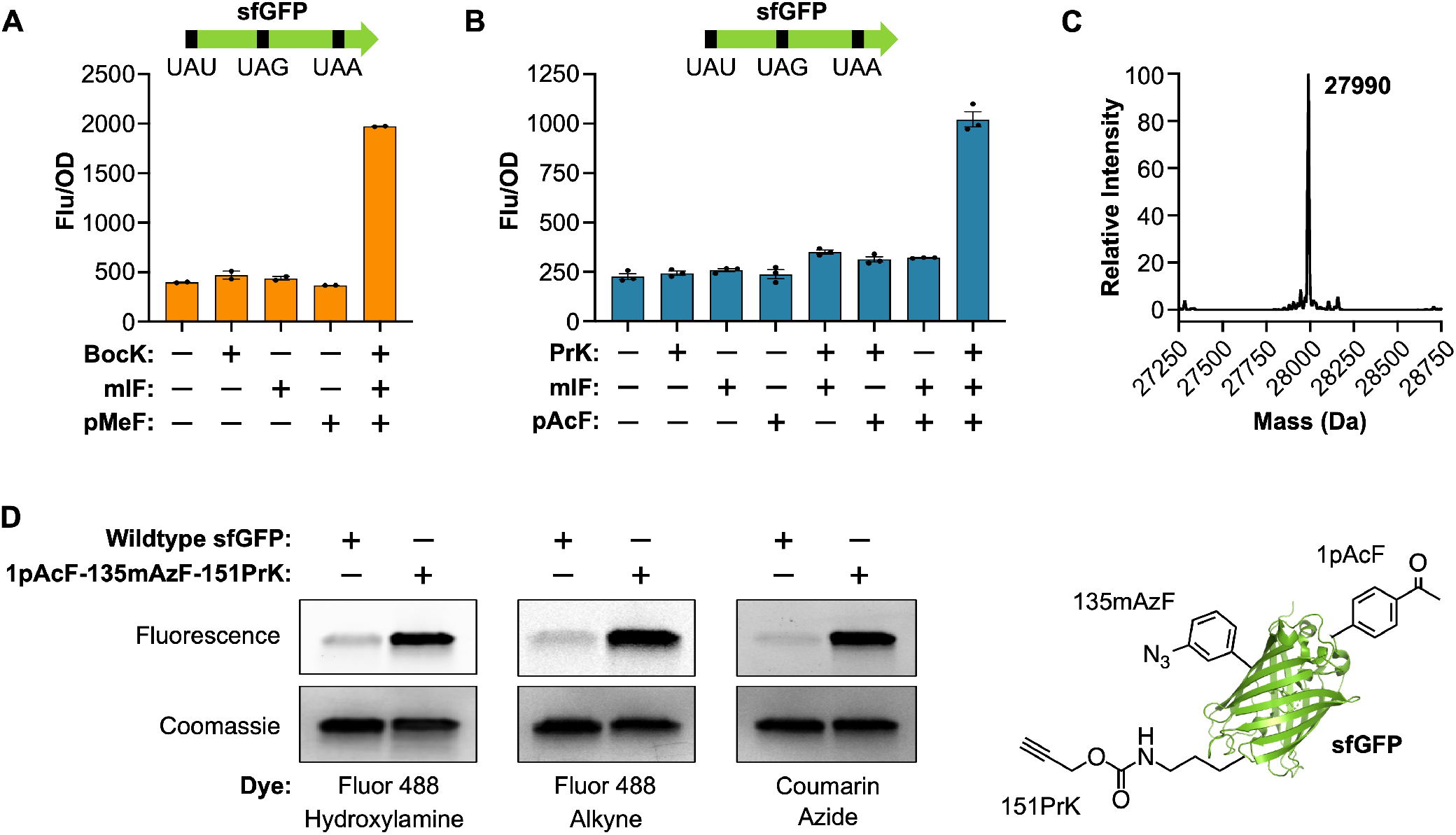
Co-translational installation of three distinct noncanonical amino acids encoded by UAU, UAG, and UAA. (A) Expression of sfGFP[1UAU-135UAG-151UAA] in the presence of BocK, mIF, and pMeF. Data are displayed as the mean ± SEM of two biological replicates. (B) Expression of sfGFP[1UAU-135UAG-151UAA] in the presence of PrK, mIF, and pAcF. Data are displayed as the mean ± SEM of three biological replicates. (C) LC-MS of sfGFP-1pAcF-135mIF-151PrK (theoretical mass = 27992 Da). (D) Labeling of sfGFP-1pAcF-135mAzF-151PrK with Fluor 488-hydroxylamine, Fluor 488-alkyne, and coumarin azide. Proteins were resolved by SDS-PAGE and visualized by in-gel fluorescence and Coomassie staining. A cartoon representation of sfGFP-1pAcF-135mAzF-151PrK is also shown (PDB: 2B3P).

In summary, we have engineered an orthogonal initiator tRNA that enables efficient reassignment of the UAU codon to initiate translation with ncAAs. We demonstrated that this system enables double and triple incorporation of distinct ncAAs encoded by UAU, UAG, and UAA codons. Several of the ncAAs that we tested for simultaneous incorporation (e.g., pAcF, mIF, mAzF, and PrK) contain unique bioorthogonal reaction handles that enable double and triple labeling of pro-teins produced from this system. In addition to the demonstrated reactivity of pAcF, mAzF, and PrK, mIF can also be used as a reaction handle for protein conjugation with boronic acids *via* palladium-catalyzed Suzuki-Miyaura cross-coupling.^49^

### Safety statement

No unexpected, new, or significant haz-ards or risks were encountered during the course of this work.

## ASSOCIATED CONTENT

### Supporting Information

The Supporting Information is available free of charge on the ACS Publications website.

Materials and Methods, Supplementary Figures and Tables, DNA Sequences (PDF)

## AUTHOR INFORMATION

### Notes

The authors declare no competing financial interests.

## ACKNOWLEDGMENT

We thank Kyle Hoffman and Sebasthian Santiago for help with mass spectrometry data collection and analysis, and Erol Vatansever, Jonathan Fischer, Christina Chung, and Natalie Krahn for providing critical feedback on the manuscript. J.M.T. was supported by the Center for Genetically Encoded Materials, an NSF Center for Chemical Innovation (NSF CHE-2021739 to A.S.), O.V.-R. was supported by the National Institute of General Medical Sciences (R35GM122560 to D.S.). The genetic experiments were supported by the Division of Chemical Sciences, Geosciences, and Biosciences, Office of Basic Energy Sciences of the Department of Energy (DE-FG0298ER20311 to D.S.).

