## Supplementary Information for "Genetic Encoding of Three Distinct Noncanonical Amino Acids Using Reprogrammed Initiator and Nonsense Codons"

**Contents**

**Materials and Methods** ... **3**
General ... 3
Safety Statement ... 3
Plasmid Construction ... 3
Protein Expression and Purification ... 5
*In vivo* sfGFP Reporter Assay ... 5
*In vitro* tRNA Aminoacylation ... 5
*Ma*PylRS N166/V168 Library Screening ... 6
Protein Labeling ... 6
Mass Spectroscopy ... 7
**Supplementary Figures** ... **8**
Figure S1. ... 8
Figure S2 ... 8
Figure S3 ... 9
Figure S4 ... 9
Figure S5 ... 10
Figure S6 ... 11
Figure S7 ... 12
Figure S8 ... 13
Figure S9 ... 14
Figure S10 ... 15
Figure S11 ... 16
Figure S12 ... 16
Figure S13 ... 17
Figure S14 ... 18
Figure S15 ... 19
Figure S16 ... 20
Figure S17 ... 21
Figure S18 ... 22
Figure S19 ... 23
Figure S20 ... 24
**Supplementary Tables** ... **25**
Table S1. Plasmids used in this study ... 25
Table S2. Oligonucleotides used in this study ... 27
**DNA Sequences** ... **29**
**References** ... **32**

#### Materials and Methods

##### General

All ncAAs were purchased from Chem-Impex International, Click Chemistry Tools, 1 ClickChemistry, or Sigma-Aldrich, and used without further purification. Enzymes and reagents for molecular cloning were purchased from New England Biolabs (NEB) and Takara Bio USA. Oligonucleotide synthesis was performed by the Keck Biotechnology Resource Laboratory at Yale University. DNA sequencing services were provided by the Keck Biotechnology Resource Laboratory and Quintara Biosciences. Mass spectrometry services were provided by Bioinformatics Solutions Inc. Unless noted, antibiotics for cell cultures were used at the following concentrations: ampicillin (Amp), 100 µg/mL; spectinomycin (Spec), 50 µg/mL; chloramphenicol (Cm), 34 µg/mL; tetracycline (Tet), 12 µg/mL.

##### Safety Statement

No unexpected, new, or significant hazards or risks were encountered during the course of this work.

##### Plasmid Construction

For a complete list of the plasmids used in this study see Table S1. Primer sequences are given in Table S2.

*pBAD-sfGFP[2UAG]-MmtRNA$^{Pyl}_{CUA}$* (pJT008) and *pBAD-sfGFP[2UAG]-MatRNA(6)$^{Pyl}_{CUA}$* (pJT010): plasmids pJT008 and pJT010 were constructed as described previously.[1] Briefly, tRNA$^{Pyl}$ expression cassettes under control of an Lpp promoter were PCR-amplified from preexisting pCAM-cat[1UAG]-tRNA$^{Pyl}$ constructs, using primers JT27/JT28. The amplified fragments were digested with *SphI* and cloned into *SphI*-digested pBAD-sfGFP[2UAG] using the In-Fusion® HD Cloning Kit (Takara Bio Inc.).

*pBAD-sfGFP[1NNN]-itRNA$^{Ty2}_{CUA}$:* (pJT100-pJT107): The initiating methionine codon of sfGFP was mutated to UAC, UAU, UUC, UGC, UCC, AAC, GAC, or CAC by amplifying the previously reported plasmid pJT052[1] with the reverse primer JT261 and forward primers JT262–JT268. The PCR products were circularized using the In-Fusion® HD Cloning Kit.

*pBAD-sfGFP[1NNN]-itRNA$^{Ty2}_{NNN}$* (pJT108-pJT115): The anticodon of *i*tRNA$^{Ty2}_{CUA}$ in plasmids pJT100-pJT107 was mutated to match the initiating codon of sfGFP[1NNN] by amplifying the plasmids with the reverse primer JT271 and the forward primers JT272–JT279. The PCR products were circularized using NEBuilder® HiFi DNA Assembly Master Mix (NEB).

*pBAD-sfGFP$^{Opt}$[1UAU]-itRNA$^{Ty2}_{AUA}$* (pJT117): A codon optimized variant of sfGFP in which all UAU codons were replaced with UAC (sfGFP$^{Opt}$) was synthesized by Integrated DNA Technologies. The synthetic gene was amplified with primers JT292/JT293 while a recipient vector (pJT109) was amplified with primers JT290/JT291. The PCR products were assembled using NEBuilder® HiFi DNA Assembly Master Mix.

*pBAD-sfGFP$^{Opt}$[1UAU-135UAG-151UAA]- itRNA$^{Ty2}_{AUA}$* (pJT121): The template plasmid pJT117 was amplified with primers JT296/JT297 to introduce a Y151→UAA mutation in the sfGFP[1UAU] gene. The resulting plasmid was subsequently amplified with primers JT294/JT295 to introduce a N135→UAG mutation affording sfGFP$^{Opt}$[1UAU-135UAG-151UAA].

*pBAD-sfGFP$^{Opt}$[1UAU-135UAG-151UAA]-itRNA$^{Ty2}_{AUA}$-MmtRNA$^{Pyl}_{AAU}$* (pJT151): The plasmid pEVOL-*Mm*tRNA$^{Pyl}_{CUA}$-*Mm*PylRS(N346A/C348A) was a gift from Professor Wenshe Liu at Texas A&M University (Addgene plasmid #127411).[2] The *Mm*tRNA$^{Pyl}_{CUA}$ expression cassette was amplified from this plasmid using the primer pair JT317/JT318. A recipient plasmid (pJT121) was simultaneously linearized with primers JT313/JT314. The purified PCR products were assembled using NEBuilder® HiFi DNA Assembly Master Mix to afford *pBAD-sfGFP$^{Opt}$[1UAU-135UAG-151UAA]-itRNA$^{Ty2}$$_{AUA}$-MmtRNA$^{Pyl}$$_{CUA}$*. This plasmid was subsequently amplified with primers JT356/JT357 to mutate the anticodon of *Mm*tRNA$^{Pyl}$ from CUA to UAA and to introduce a previously reported U:G→C:G mutation in the anticodon stem for improved suppression efficiency.[3]

*pBAD-sfGFP$^{Opt}$[1UAU-135UAG-151UAA]-itRNA$^{Ty2}$$_{AUA}$-MatRNA(6)$^{Pyl}$$_{CUA}$* (pJT166): The *MatRNA(6)$^{Pyl}$$_{CUA}$* expression cassette under control of a the *proK* promoter was PCR-amplified from plasmid pJT159 using primers JT396/JT397. The recipient plasmid (pJT121) was simultaneously linearized with primers JT394/JT395. The purified PCR products were then assembled with NEBuilder® HiFi DNA Assembly Master Mix.

*pBAD-sfGFP$^{Opt}$[1UAU-135UAG]-itRNA$^{Ty2}$$_{AUA}$-MatRNA(6)$^{Pyl}$$_{CUA}$-MmtRNA$^{Py}$$_{UUA}$* (pJT171): Plasmid pJT167 was constructed in the same manner as pJT166 but using pJT151 as the recipient plasmid. pJT167 was amplified with primers JT362/JT363 to revert the 151UAA mutation, affording pJT171.

*pBAD-sfGFP$^{Opt}$[1UAU-151UAA]-itRNA$^{Ty2}$$_{AUA}$-MatRNA(6)$^{Pyl}$$_{CUA}$-MmtRNA$^{Pyl}$$_{UUA}$* (pJT172): Like pJT171, pJT172 was constructed using pJT167 as a template. pJT167 was amplified with primers JT360/JT361 to revert the 135UAG mutation affording pJT172.

*pBAD-sfGFP$^{Opt}$[1UAG]-itRNA$^{Ty2}$$_{AUA}$* (pJT122): The synthetic sfGFP$^{Opt}$ gene was amplified with primers JT293/JT302. The recipient vector (pJT052) was simultaneously amplified with primers JT290/JT291. The PCR products were then assembled using NEBuilder® HiFi DNA Assembly Master Mix.

*pBAD-sfGFP$^{Opt}$[2UAG]-itRNA$^{Ty2}$$_{AUA}$* (pJT123): The plasmid pJT117 was amplified with primers JT300/JT301 to revert the 1UAU mutation and introduce an 2UAG mutation. The PCR product was then assembled using NEBuilder® HiFi DNA Assembly Master Mix.

*pBAD-sfGFP$^{Opt}$[2UAA]-itRNA$^{Ty2}$$_{AUA}$* (pJT124): The plasmid pJT117 was amplified with primers JT298/JT299 to revert the 1UAU mutation and simultaneously introduce an 2UAA mutation. The PCR product was then assembled using NEBuilder® HiFi DNA Assembly Master Mix.

*pULTRA-MmPylRS-MmtRNA$^{Pyl}$$_{UUA}$* (pJT156): Wildtype *Mm*PylRS was amplified from a preexisting pEVOL plasmid using primers JT348/JT349. A previously reported recipient plasmid (pJT079) was simultaneously linearized with primers JT347/JT169.[1] The purified PCR products were then assembled with NEBuilder® HiFi DNA Assembly Master Mix to afford pULTRA-*Mm*PylRS-*MatRNA$^{Pyl}$$_{UUA}$*. The *MatRNA$^{Pyl}$$_{UUA}$* in this plasmid was replaced with *MmtRNA$^{Pyl}$$_{UUA}$* by amplifying the plasmid with primers JT356/JT357.

*pULTRA-MaPylRS-MatRNA(6)$^{Pyl}$$_{CUA}$* (pJT173): The anticodon of *MatRNA(6)$^{Pyl}$$_{UUA}$* in the previously reporter plasmid pJT079[1] was mutated to CUA by amplifying the plasmid with primers JT355/JT176 to afford pJT173.

*pSTART-AzFRS.2.t1-MaPylRS(N166S)-MatRNA(6)$^{Pyl}$$_{CUA}$* (pJT159 and pJT160): The plasmid pEVOL-AzFRS.2.t1-*Mj*tRNA$^{Tyr}$$_{CUA}$ was a kind gift from Professor Farren Isaacs at Yale University (Addgene plasmid #73546). To clone *Ma*PylRS into the preexisting *glnS* promoter, this plasmid was amplified with primers JT351/JT352. Simultaneously, *Ma*PylRS(N166S) was amplified from pJT138 using primers JT353/354. The two PCR products were then assembled using NEBuilder® HiFi DNA Assembly Master

Mix. *Mj*tRNA$^{Tyr}_{CUA}$ was then replaced with *Ma*tRNA(6)$^{Pyl}_{CUA}$ by amplifying the resultant plasmid with JT176/JT355 to afford pEVOL-AzFRS.2.t1-*Ma*PylRS(N166S)-*Ma*tRNA(6)$^{Pyl}_{CUA}$. The glnS-*Ma*PylRS(N166S)-proK-*Ma*tRNA(6)$^{Pyl}_{CUA}$-Cm$^R$ portion of this plasmid was amplified with primers JT372/JT373 and cloned into pMW-AzFRS.2.t1 (linearized with primers JT370/371) to afford pJT159. To change the *glnS* promoter for *Ma*PylRS(N166S) to the stronger lac promoter, pJT159 was amplified with primers JT386/JT387 and lac-*Ma*PylRS(N166S) was simultaneously amplified from pJT138 using primers JT388/389. The PCR products were assembled with NEBuilder® HiFi DNA Assembly Master Mix to afford pJT160.

*pSTART-AzFRS.2.t1* (pJT174): To remove *Ma*PylRS(N166S)- *Ma*tRNA(6)$^{Pyl}_{CUA}$ from pJT159 the plasmid was amplified with primers JT398/JT399 and assembled with NEBuilder® HiFi DNA Assembly Master Mix to afford pJT174.

**Protein Expression and Purification**

All sfGFP mutants were expressed in the *E. coli* strain DH10BΔ*metZWV*. For a typical experiment, freshly transformed colonies of DH10BΔ*metZWV* containing the appropriate plasmids (Table S1) were grown overnight in 2xYT media supplemented with antibiotics. The following day, overnight cultures were diluted 1:100 (1:20 for three-plasmid systems) in chemically defined media[1] and grown at 37°C to an $OD_{600}$ of 0.25-0.5, at which point protein expression was induced with the addition of 1 mM IPTG, 0.2% arabinose, and 2 mM of the indicated ncAA(s). Cells were cultured for an additional 18-20 hours and then harvested by centrifugation.

Cells were lysed by resuspending pellets in BugBuster® 10x Protein Extraction Reagent (Millipore-Sigma) that was diluted in lysis buffer (50 mM Tris pH 8.0, 500 mM NaCl, 10 mM imidazole) and supplemented with 25 U/mL Benzonase® Nuclease (Sigma). Following lysis, the solution was clarified by centrifugation (10,000 × *g*, 30 min) and proteins were purified in a gravity flow column using TALON® Metal Affinity Resin (Clonetech). The resin was washed with lysis buffer and bound proteins were eluted with elution buffer (50 mM Tris pH 8.0, 100 mM NaCl, 250 mM imidazole). Purified proteins were concentrated, and the buffer was exchanged to 25 mM sodium phosphate pH 7.4, 25 mM NaCl using Amicon® Ultra Centrifugal Filters (10 kDa NMWL). Final protein concentrations were estimated using the absorbance at 280 nm.

***In vivo* sfGFP Reporter Assay**

For a typical experiment, freshly transformed colonies of *E. coli* DH10B or DH10BΔ*metZWV* (as indicated), containing the appropriate plasmids (Table S1), were isolated and grown to saturation in 2xYT supplemented with antibiotics. Saturated cultures (5 µL) were used to inoculate 150 µL of chemically defined media[1], supplemented with antibiotics, 0.1 mM IPTG, 0.2% arabinose, and 2 mM of the indicated ncAA(s), in black, clear bottom, 96-well plates. Cultures were grown in a BioTek Synergy microplate reader at 37°C with 12 min of continuous shaking every 15 min. Fluorescence intensity ($\lambda_{ex}$ = 485 nm, $\lambda_{em}$ = 528 nm) and $OD_{600}$ were measured every 15 min for the duration of the experiment. Unless noted otherwise, data are reported as the fluorescence intensity divided by the $OD_{600}$ at the 18-hour timepoint.

***In vitro* tRNA Aminoacylation**

*Preparation of tRNA Transcripts*: The genes for *Ec*tRNA$^{Tyr}$ and *i*tRNA$^{Ty2}_{AUA}$ were cloned into pUC18 under the T7 promoter using NEBuilder® HiFi cloning. Briefly, primers 418.F and 418.R were used to amplify pUC18 for cloning of *i*tRNA$^{Ty2}_{AUA}$. 417.F and 417.R primers, containing the *i*tRNA$^{Ty2}_{AUA}$ gene were annealed prior to ligation with linearized pUC18. Similarly, primers 407.F and 407.R were used to amplify pUC18 for *Ec*tRNA$^{Tyr}$ cloning, while 413.F and 413.R primers (containing the *Ec*tRNA$^{Tyr}$ gene) were annealed before ligation into pUC18. *In vitro* transcription of *Ec*tRNA$^{Tyr}$ and *i*tRNA$^{Ty2}_{AUA}$ was carried out using DNA fragments from the pUC18 constructs amplified with the pUC18_un primer together with the

436 and 437 primers for *i*tRNA$^{Ty2}_{AUA}$ and *Ec*tRNA$^{Tyr}$, respectively. T7 RNA polymerase was purified following a previously published protocol.[4] The *in vitro* run-off transcription was performed in 40 mM Tris-HCl pH 8, 1 mM spermidine, 0.01% Triton X-100, 0.005 mg/mL BSA, 10 mM DTT, 20 mM $MgCl_2$, 3 mM NTPs, and 10 µg DNA template for 7 h at 37ºC.

*Preparation of E. coli TyrRS*: His-tagged *E. coli* TyrRS (*Ec*TyrRS) was purified using an *E. coli* strain from the ASKA collection harboring the *Ec*TyrRS expression plasmid.[5] Cells were grown to an $OD_{600}$ of 0.6 followed by induction of *Ec*TyrRS expression with 1 mM IPTG for 4 h at 37ºC. Cells were collected by centrifugation and resuspended in buffer containing 50 mM Tris pH 8, 300 mM NaCl, and protease inhibitor tablets (Roche). Cell lysis was carried out using lysozyme followed by sonication. Cell lysate was cleared by centrifugation at 19000 × *g* for 50 min at 4ºC. The His-tagged protein was isolated using TALON® Metal Affinity Resin (Clonetech) and eluted using varying concentration of imidazole. The purified protein was concentrated and stored at -20ºC in buffer containing 12.5 mM HEPES pH 7.3, 75 mM NaCl, and 40% glycerol. Protein concentration was determined using the Bradford assay.[6]

*Aminoacylation Reactions*: Aminoacylation reactions were performed at 37ºC in a solution containing 50 mM HEPES pH 7.3, 4 mM ATP, 10 mM $MgCl_2$, 3 mM KCl, 2.5 µM [$^3$H]-tyrosine (PerkinElmer), 5 µM tRNA, and 0.5 µM *Ec*TyrRS. At indicated time points, 10 µL of reaction mixture was quenched on a filter pad (Whatman 3MM) pre-soaked with 5% TCA. Pads were washed three times with 5% TCA followed by a final wash with absolute ethanol. Pads were air-dried and radioactivity was measured using a scintillation counter. A reaction in the absence of tRNA was used to measure the background level of Tyr binding to the filter in the presence of *Ec*TyrRS. All data points are the average of at least three independent experiments with the standard deviation indicated. Data were plotted using the GraphPad Prism software.

**_Ma_PylRS N166/V168 Library Screening**

The two-site *Ma*PylRS library was prepared by amplifying the plasmid pJT065 (containing the wildtype *Ma*PylRS gene) with primers JT269/JT270. The PCR product was digested with *DpnI* and then circularized using NEBuilder® HiFi DNA Assembly Master Mix. The cloned product was then used to transform *E. coli* DH10B containing the selection plasmid pCAM-Ma (Table S1). Transformed cells were recovered at 37ºC for 1 hour and then plated on chemically defined agar plates[1] supplemented with Spec, Tet, 50 µg/mL Cm, 1 mM IPTG, and 1 mM mIF or oMeF. Plates were incubated at 37ºC for ~20 hours and then surviving clones were screened for ncAA recognition by replating on plates with and without an ncAA (Figure S11).

**Protein Labeling**

*3-azido-7-hydroxycoumarin and Fluor 488-alkyne Labeling:* To 92 µL of reaction buffer (25 mM sodium phosphate pH 7.4, 25 mM NaCl) was added $CuSO_4$ (2 µL, 5 mM) and BTTAA (2 µL, 10 mM in DMSO, Click Chemistry Tools). The solution was mixed and then added to a sample of protein (100 µL, 20 µM) in reaction buffer. To the resulting mixture was added, sequentially, 3-azido-7-hydroxycoumarin (2 µL, 10 mM in DMSO, Santa Cruz Biotechnology) or fluor 488-alkyne (2 µL, 10 mM in DMSO, Sigma-Aldrich) and sodium ascorbate (2 µL, 250 mM). The resulting solution was incubated at room temperature for 2 hours. After the reaction, excess dye and reagents were removed by three rounds of 5-fold dilution in reaction buffer, followed by concentration using Amicon® Ultra Centrifugal Filters (3 kDa NMWL).

*HiLyte™ Fluor 488-hydroxylamine Labeling:* Protein samples (25 µL, 40 µM in 25 mM sodium phosphate pH 7.4, 25 mM NaCl) were diluted with 75 µL of a low pH buffer (100 mM potassium phosphate pH 4.5, 250 mM NaCl) resulting in a final solution pH of ~5.8. To the diluted samples was added HiLyte™ Fluor 488-hydroxylamine (2 µL, 25 mM in DMSO, AnaSpec Inc.). The solution was mixed and then incubated at 25ºC for 18 hours. After the reaction, excess dye was removed by three rounds of 5-fold dilution in reaction buffer, followed by concentration using Amicon® Ultra Centrifugal Filters (3 kDa NMWL).

**Mass Spectroscopy**

The mass spectroscopy data shown in Figure 1E were collected exactly as reported previously.[1] All other LC-MS and MS/MS were performed by Bioinformatics Solutions Inc. in Waterloo, Ontario, Canada as follows:

*Intact Protein LC-MS*: LC-MS analysis of intact proteins was performed on a Thermo Scientific Orbitrap Fusion Lumos Tribrid mass spectrometer equipped with a heated electrospray ionization source with a Thermo Fisher Ultimate 3000 RSLCnano HPLC System. Proteins were analyzed on a MAbPac RP analytical column (4 μm, 3.0 × 50 mm, 70°C, ThermoFisher) using a mobile phase consisting of water/acetonitrile in 0.1% formic acid. Proteins were separated at a rate of 500 μL/min under the following gradient: 0–11 min, 10–45% acetonitrile; 11–13 min, 45–95% acetonitrile; 13–15 min, 95% acetonitrile; 15–17 min, 20–10% acetonitrile; 17–20 min, 10% acetonitrile. Data were collected using full scans at 15000 resolution in the orbitrap over an *m/z* range of 700-2200 in positive ion mode. The maximum injection time was limited to 50 ms, with an AGC target of 4e6. Ten micro scans were employed with the RF lens set to 45%. 15 V of insource CID was applied.

*LC-MS/MS*: Reduced protein samples were alkylated with iodoacetamide prior to digestion with trypsin or chymotrypsin. LC-MS/MS analysis was performed on a Thermo Scientific Orbitrap Fusion Lumos Tribrid mass spectrometer, equipped with a nanospray ionization source, in positive ion mode, and a Thermo Fisher Ultimate 3000 RSLCnano HPLC System. Digested proteins were loaded on a PepMap$^{TM}$ 100 C18 trap column (5 μm, 60°C, ThermoFisher) with a flow of 30 μL/min. Peptides were eluted at a rate of 0.2 μL/min and separated on a ReproSil C18 analytical column (1.9 μm, PepSep) using a water/acetonitrile mobile phase, in 0.1% formic acid, and over the following gradient: 0–45 min, 4–35% acetonitrile; 45–55 min, 90% acetonitrile; 55–60 min, 4% acetonitrile. Data were collected using data-dependent mode with a cycle time of 3 seconds. MS1 scan was performed in orbitrap with an *m/z* range of 400–1600 and at a resolution of 120000 *m/z*. Maximum injection time was limited to 50 ms with an AGC target of 4e5. RF lens was set to 30%. Isolation for MS2 scans was performed in the quadrupole, with an isolation window of 0.7. MS2 scans were done in the linear ion trap at turbo scan rate, with a maximum injection time of 35 ms and AGC target at 1e4. CID was used for generating MS2 spectrum, as fixed normalized collision energy of 30% and activation time at 10 ms.

*Data Analysis*: Raw data files were processed using PEAKS XPro (v10.6, Bioinformatics Solutions Inc., Ontario, Canada). The data were searched against a custom database containing sfGFP variants with alanine substitutions where ncAA residues are inserted, in conjunction with the *E. coli* K12 Uniprot reviewed database. Parent mass tolerance was set to 10 ppm, with fragment mass tolerance of 0.6 Da. Semi-specific cleavage with trypsin or chymotrypsin was selected with a maximum of 3 missed cleavages. Fixed modifications of carbamidomethylation (57.02 Da) on cysteine residues were specified. Variable modifications of deamination (0.98 Da) on asparagine and glutamine, as well as oxidation (15.99 Da) on methionine were specified. In addition, variable modifications at alanine residues of +118.05 Da, +139.07 Da, and +201.94 Da were set for detecting insertion of pAcF, PrK, and mIF, respectively. Only peptides above a -10logP score of 22.2 and a 1% FDR were used from the PEAKS database search.

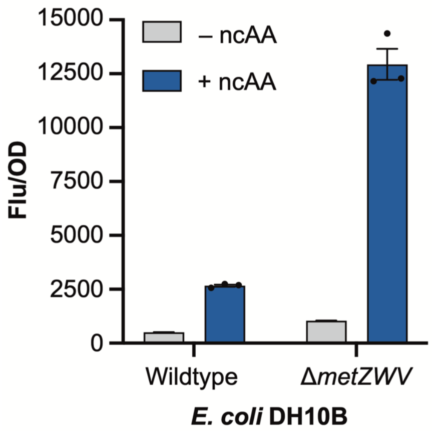


**Figure S1. A comparison of sfGFP[1UAU] expression in wildtype *E. coli* DH10B and in DH10BΔ*metZWV*.** Initiation at UAU is more efficient in DH10BΔ*metZWV*. Cells were grown in defined media[1] supplemented with 0.1 mM IPTG, 0.2% arabinose, and ± 2 mM pMeF. Data were collected 16 hours after induction and are displayed as the mean ± SEM of three biological replicates.

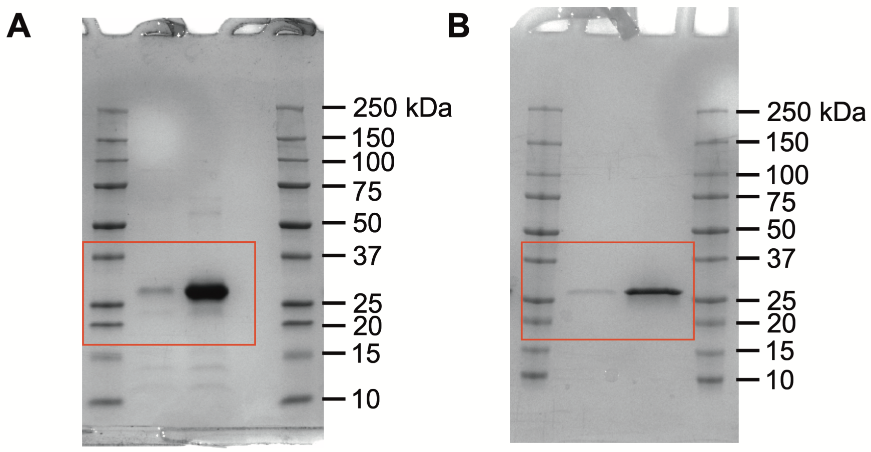


**Figure S2. Extended data for Figure 2.** Uncropped gel images for Figure 2D (A) and Figure 2G (B).

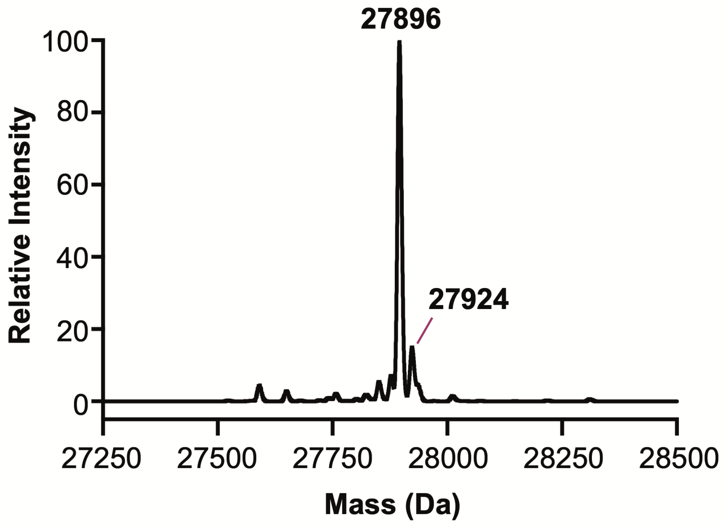


**Figure S3. LC-MS of sfGFP[1UAU] expressed with pIF.** Major peaks correspond to sfGFP with pIF incorporated at the initiating position (sfGFP-1pIF; theoretical mass = 27898 Da) and *N*-formyl-sfGFP-1pIF (theoretical mass = 27926 Da). Peaks corresponding to sfGFP with pIF incorporated at one or more elongating UAU codons were not observed (theoretical masses: pIF at 1 elongating codon = 28008 Da; 2 elongating codons = 28118 Da; 3 elongating codons = 28228 Da; 4 elongating codons = 28338 Da, 5 elongating codons = 28448 Da).

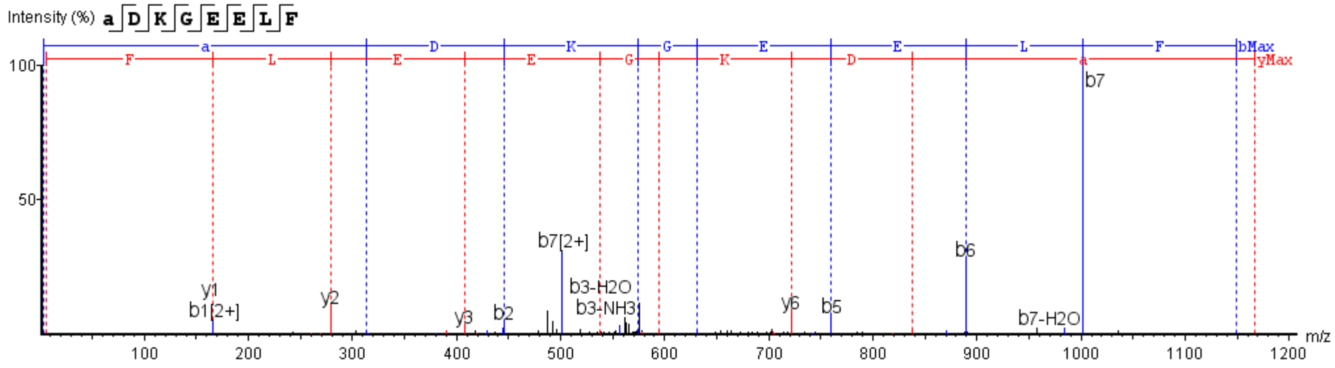


**Figure S4. MS/MS analysis of sfGFP-1pIF.** The fragmentation pattern supports incorporation of pIF at the initiating position. In the peptide sequence pIF is abbreviated with the letter *a*.

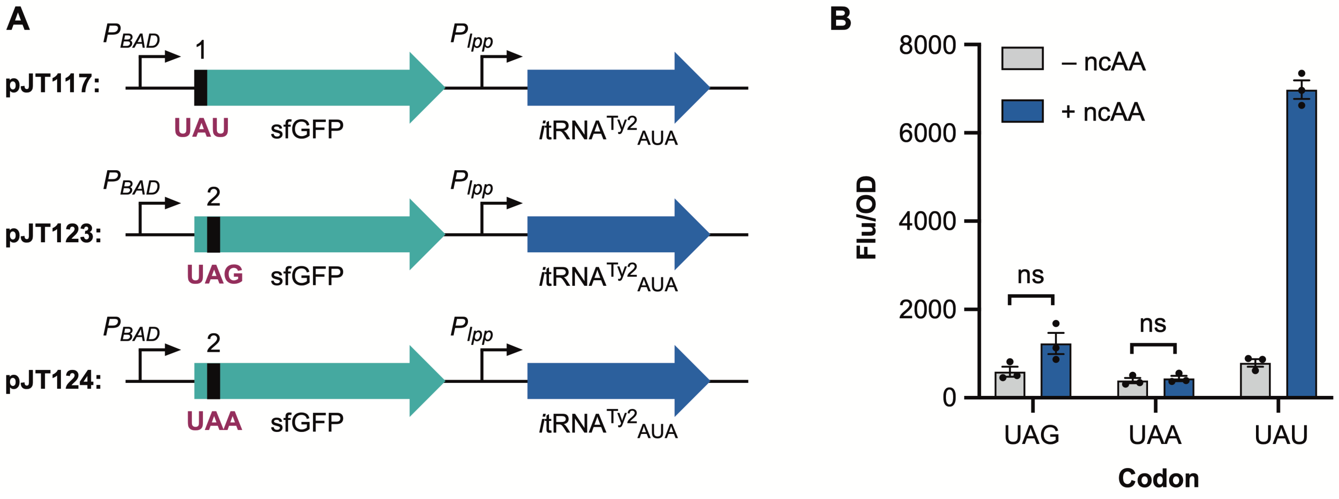


**Figure S5. Suppression of UAU, UAG, and UAA codons by *i*tRNA$^{Ty2}_{AUA}$.** *i*tRNA$^{Ty2}_{AUA}$ does not significantly suppress UAG or UAA. (A) Maps of the three reporter plasmids (pJT117, pJT123, and pJT124) used to measure expression of sfGFP with an initiating UAU codon or elongating UAG and UAA codons. (B) Flu/OD data for sfGFP[1UAU], sfGFP[2UAG], and sfGFP[2UAA] co-expressed with *i*tRNA$^{Ty2}_{AUA}$ and AzFRS.2.t1. Cells were grown in defined media[1] supplemented with 0.1 mM IPTG, 0.2% arabinose, and ± 2 mM pMeF. Data were collected 15 hours after induction and are displayed as the mean ± SEM of three biological replicates. ns = not significant (paired t-test).

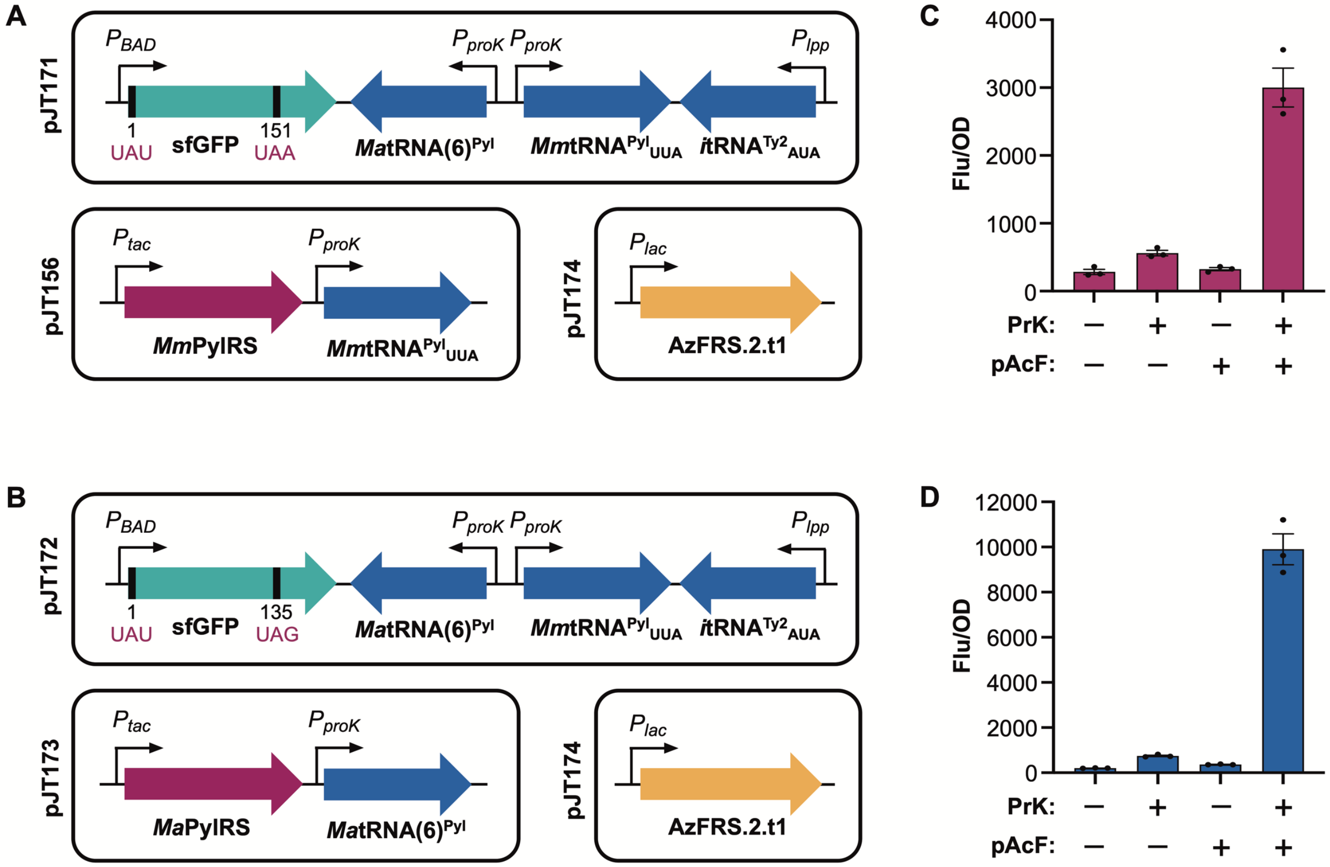


**Figure S6. Co-translational installation of two distinct noncanonical amino acids using UAU and UAG/UAA codons**. (A) Maps of the three-plasmid system used to install two ncAAs in response to an initiating UAU and elongating UAA. (B) Maps of the three-plasmid system used to install two ncAAs in response to an initiating UAU and elongating UAG. (C) Expression of sfGFP[1UAU-151UAA] in the presence pAcF and PrK. (D) Expression of sfGFP[1UAU-135UAG] in the presence of pAcF and PrK. Cells were grown in defined media[1] supplemented with 0.1 mM IPTG, 0.2% arabinose, and ± 2 mM pAcF and PrK. Data were collected 18 hours after induction and are displayed as the mean ± SEM of three biological replicates.

A

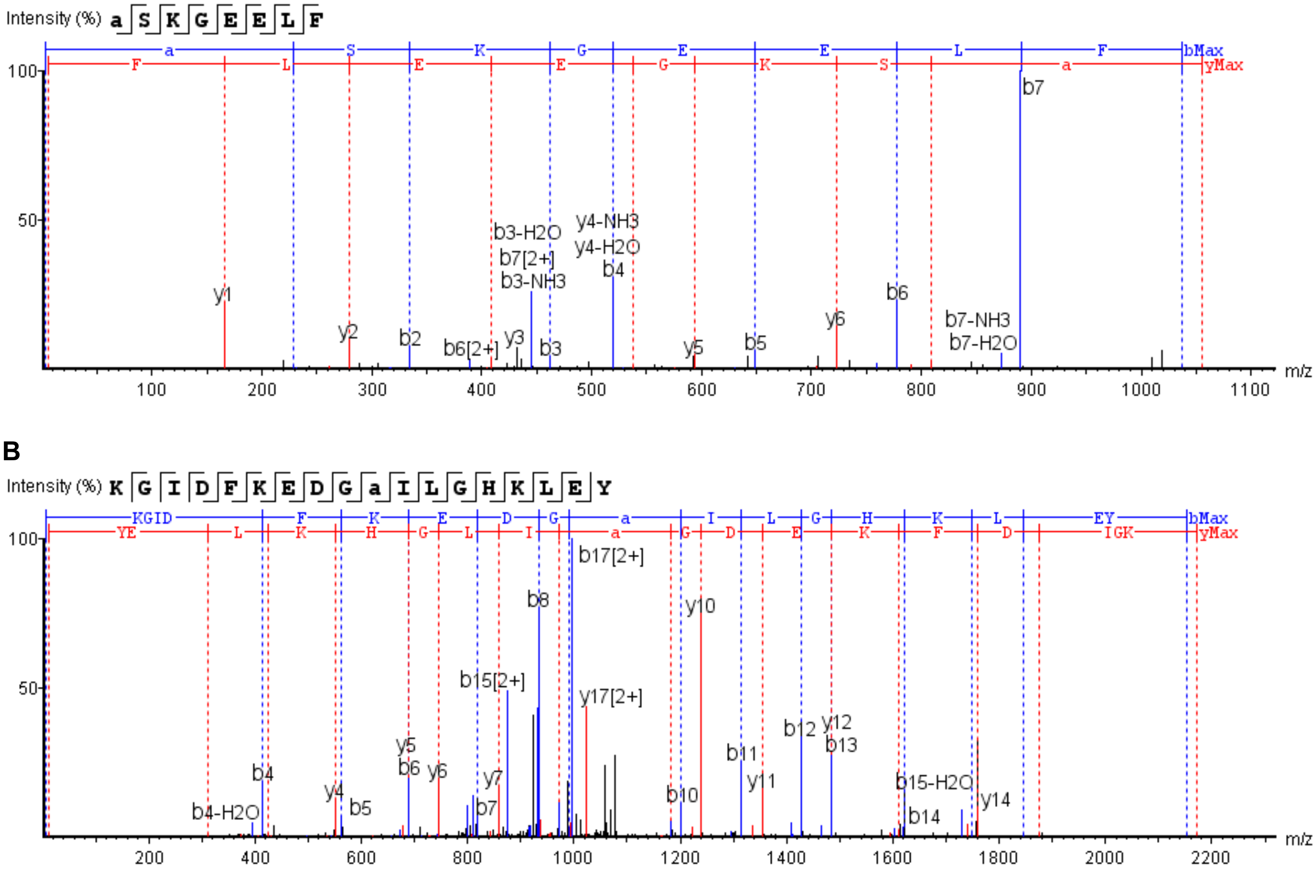


**Figure S7. MS/MS analysis of sfGFP-1pAcF-135PrK.** The spectra support the incorporation of pAcF at position 1 (A) and PrK at position 135 (B). ncAAs are abbreviated in the peptide sequence with the letter *a*.

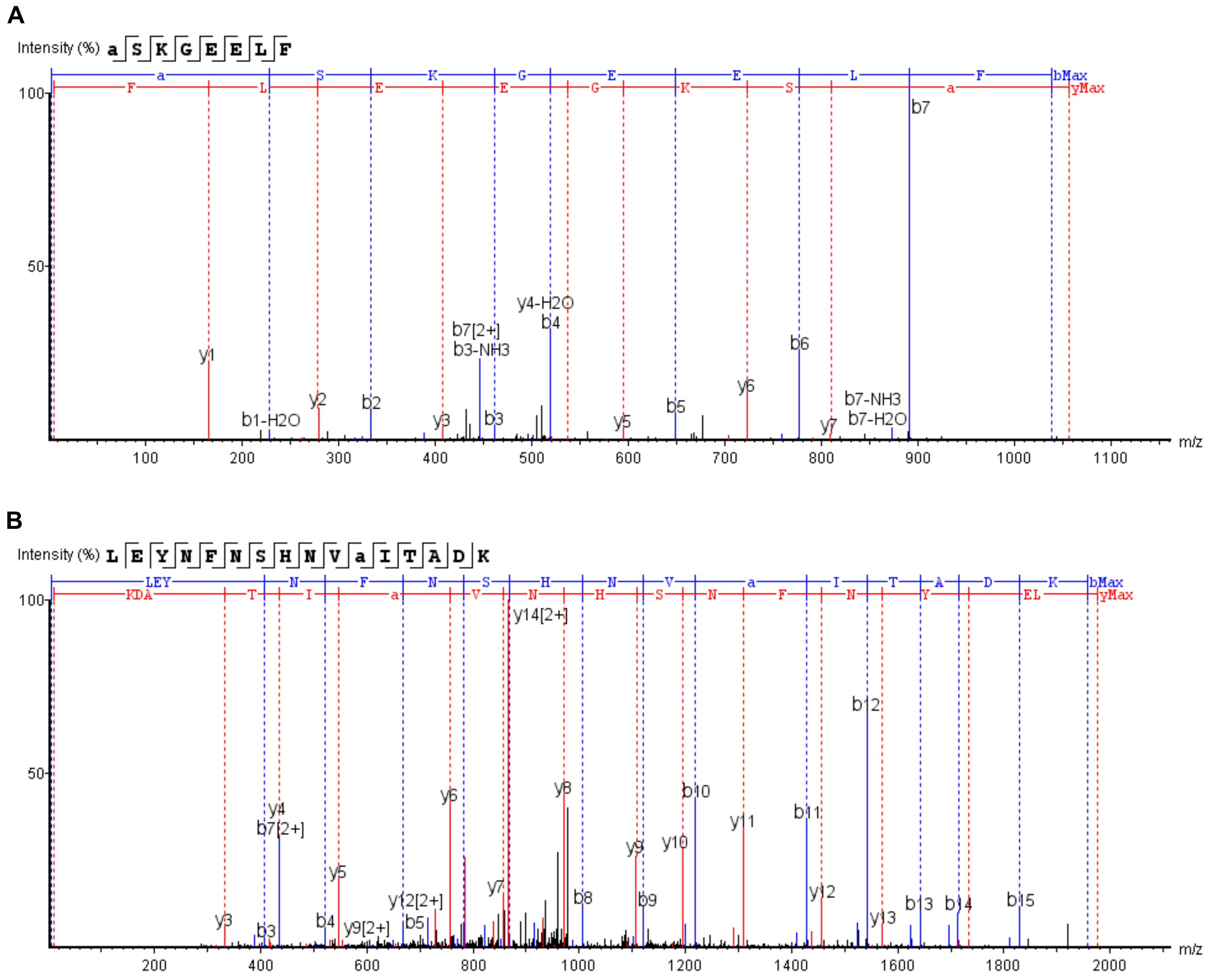


**Figure S8. MS/MS analysis of sfGFP-1pAcF-151PrK.** The spectra support the incorporation of pAcF at position 1 (A) and PrK at position 151 (B). ncAAs are abbreviated in the peptide sequence with the letter *a*.

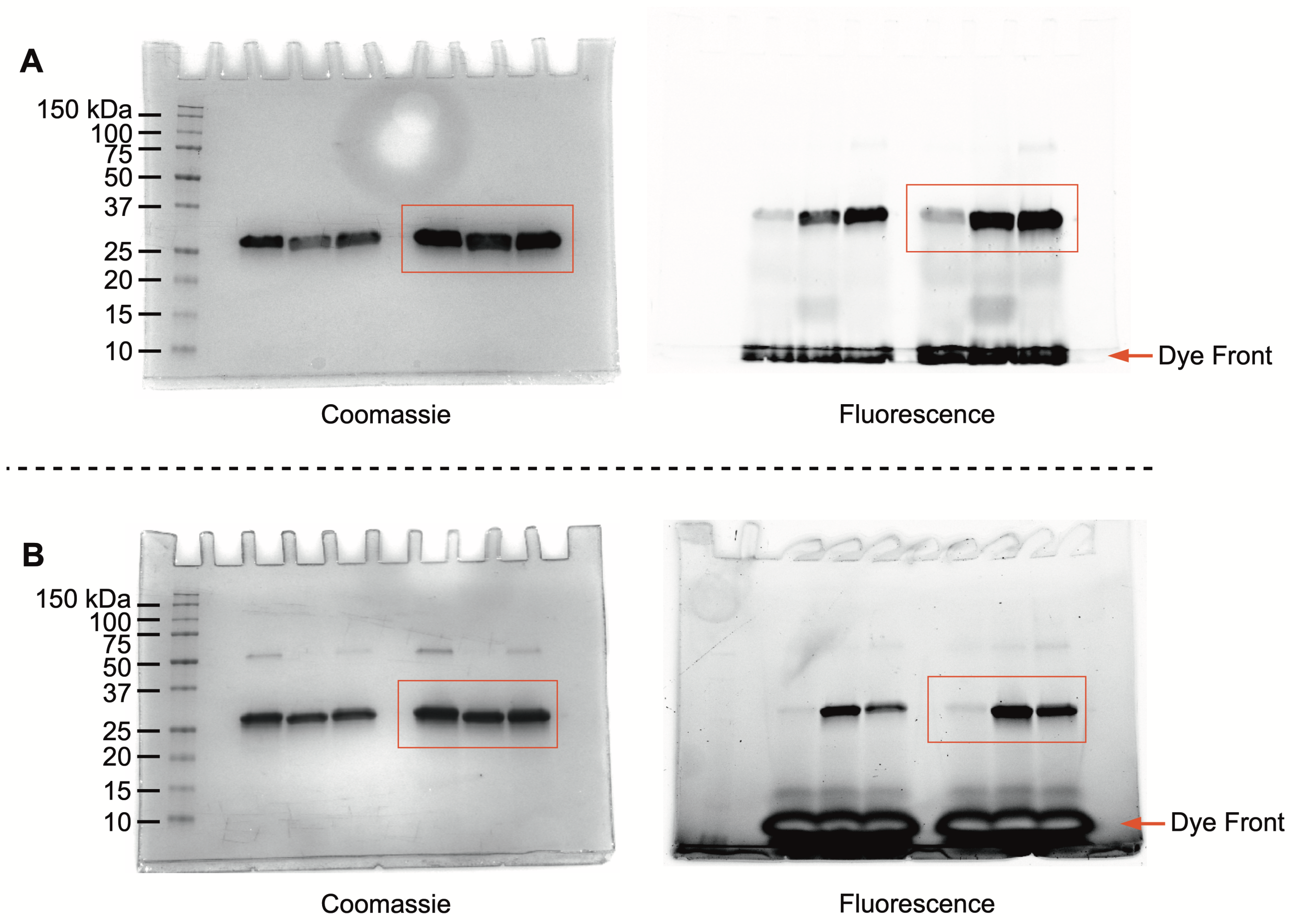


**Figure S9. Extended data for Figure 3E.** Uncropped gel images for labeling of sfGFP-1pAcF-135PrK and sfGFP-1pAcF-151PrK with Fluor 488-hydroxylamine (A) and coumarin azide (B).

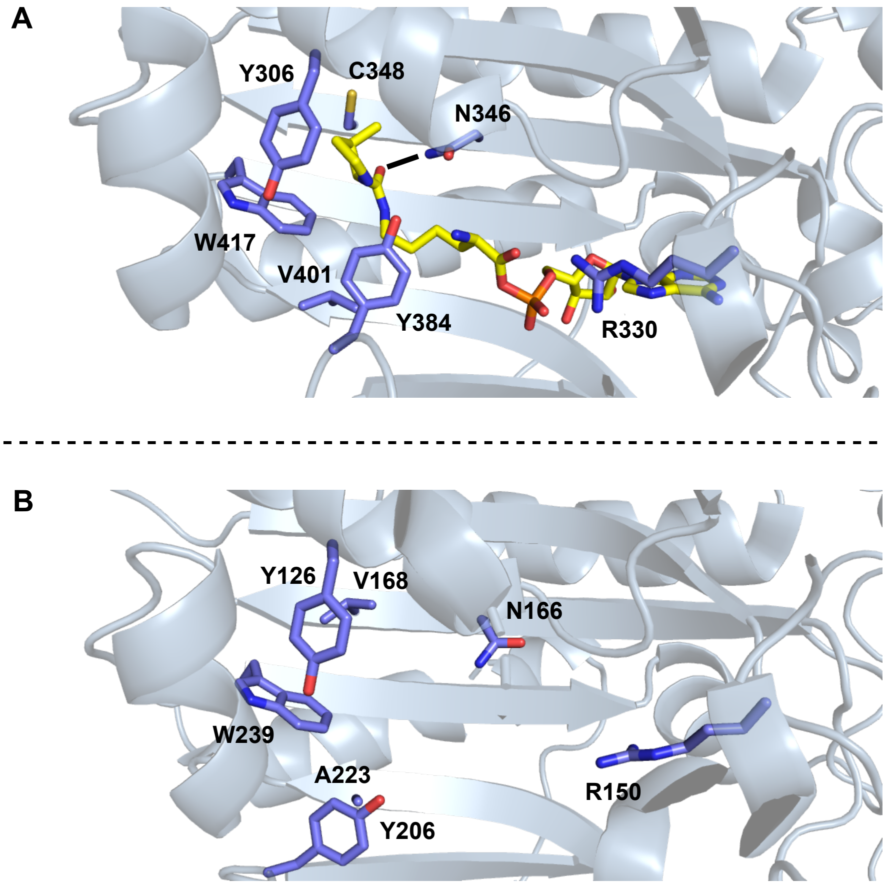


**Figure S10. A comparison of the substrate binding pockets of *Mm*PylRS and *Ma*PylRS.** Key active site residues are labeled. (A) The crystal structure of *Mm*PylRS in complex with adenylated pyrrolysine and pyrophosphate (PDB: 2Q7H). The hydrogen bond between N346 and the side chain amide of pyrrolysine is shown as a solid black line. (B) The crystal structure of *Ma*PylRS apoenzyme (PDB: 6EZD).

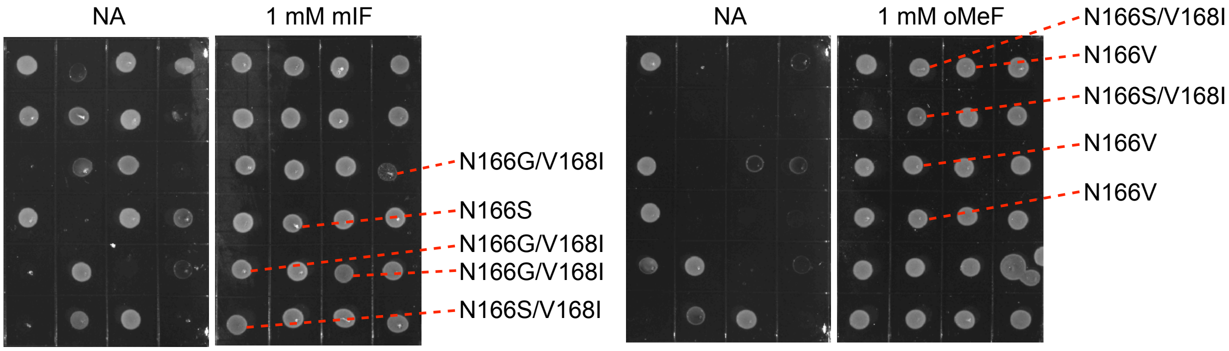


**Figure S11. *Ma*PylRS N166/V168 library screening using chloramphenicol acetyltransferase mutant cat[112UAG].** Following the first round of selection, surviving colonies were re-plated on media with and without the indicated ncAA. Clones that survived on chloramphenicol only in the presence of the ncAA were arbitrarily chosen for sequencing of the *Ma*PylRS gene. Identified mutants are labeled. Cells were grown on defined media agar[1] supplemented with 50 µg/mL chloramphenicol, spectinomycin, tetracycline, 1 mM IPTG, and 1 mM mIF, 1 mM oMeF, or no ncAA (NA).

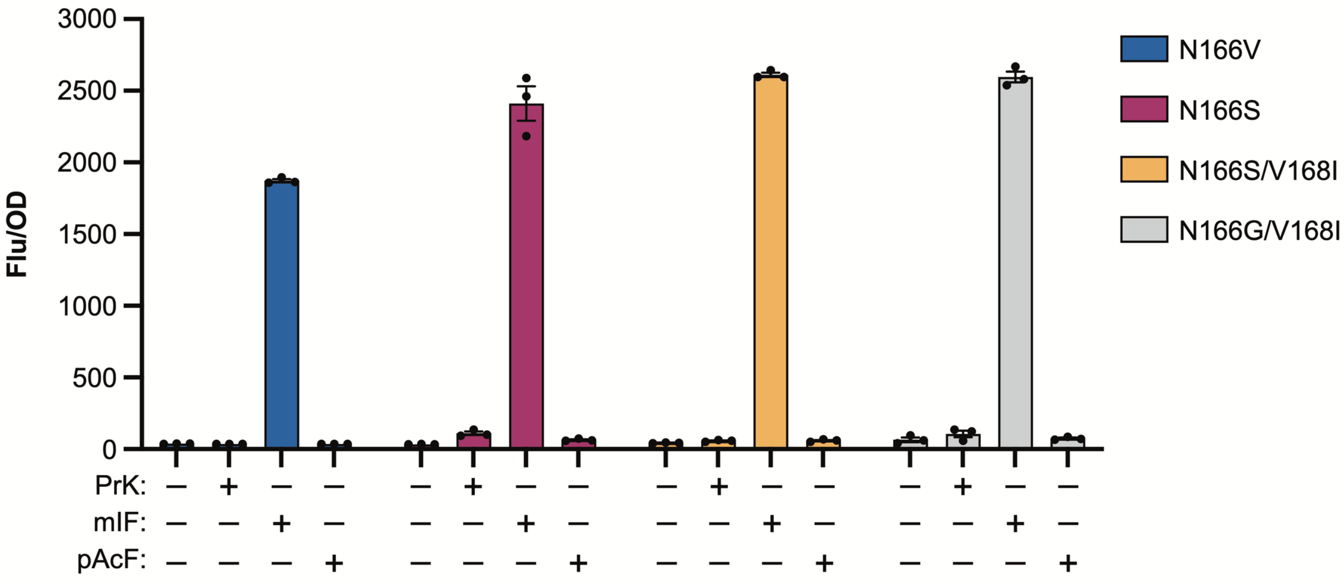


**Figure S12. Substrate specificities of four *Ma*PylRS mutants.** All mutants were selective for mIF. Substrate specificity was determined by measuring the expression of sfGFP[2UAG] in DH10B cells that are concurrently expressing *Ma*tRNA(6)$^{Pyl}$ and the indicated *Ma*PylRS mutant. Cells were grown in defined media[1] supplemented with 0.1 mM IPTG, 0.2% arabinose, and 2 mM of PrK, mIF, pAcF, or no ncAA (NA). Data were collected 18 hours after induction and are displayed as the mean ± SEM of three biological replicates.

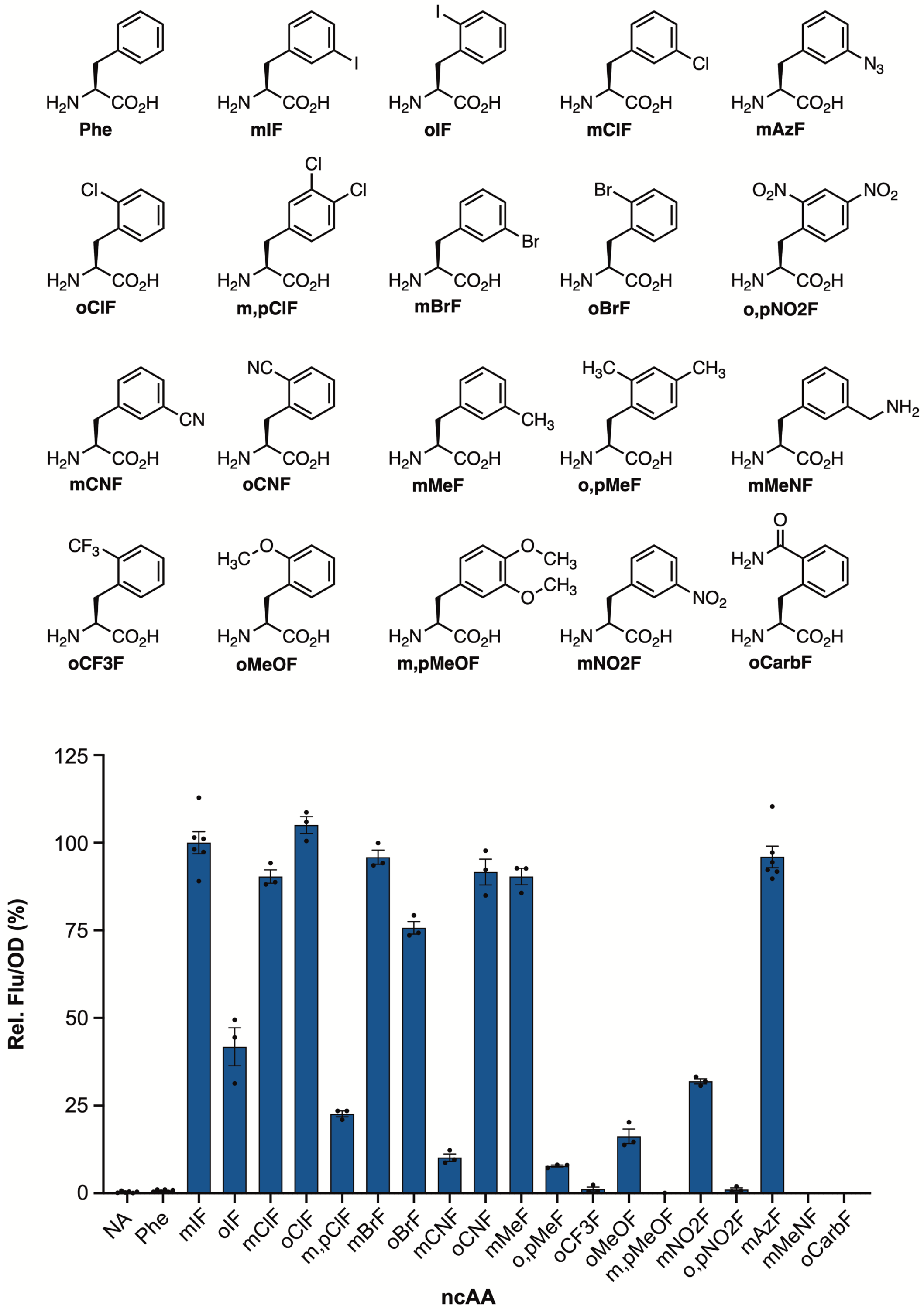


**Figure S13. Substrate specificity of *Ma*PylRS variant N166S.** (Top) Structures of amino acid with which we tested *Ma*PylRS(N166S) activity. (Bottom) Expression of sfGFP[2UAG] in DH10B cells co-expressing *Ma*PylRS(N166S) and *Ma*tRNA(6)$^{Pyl}$. Cells were grown in defined media supplemented with 0.1 mM IPTG, 0.2% arabinose, and 2 mM of the indicated amino acid. Data were collected 18 hours after induction and are displayed as the mean ± SEM of at least three biological replicates. NA = not added.

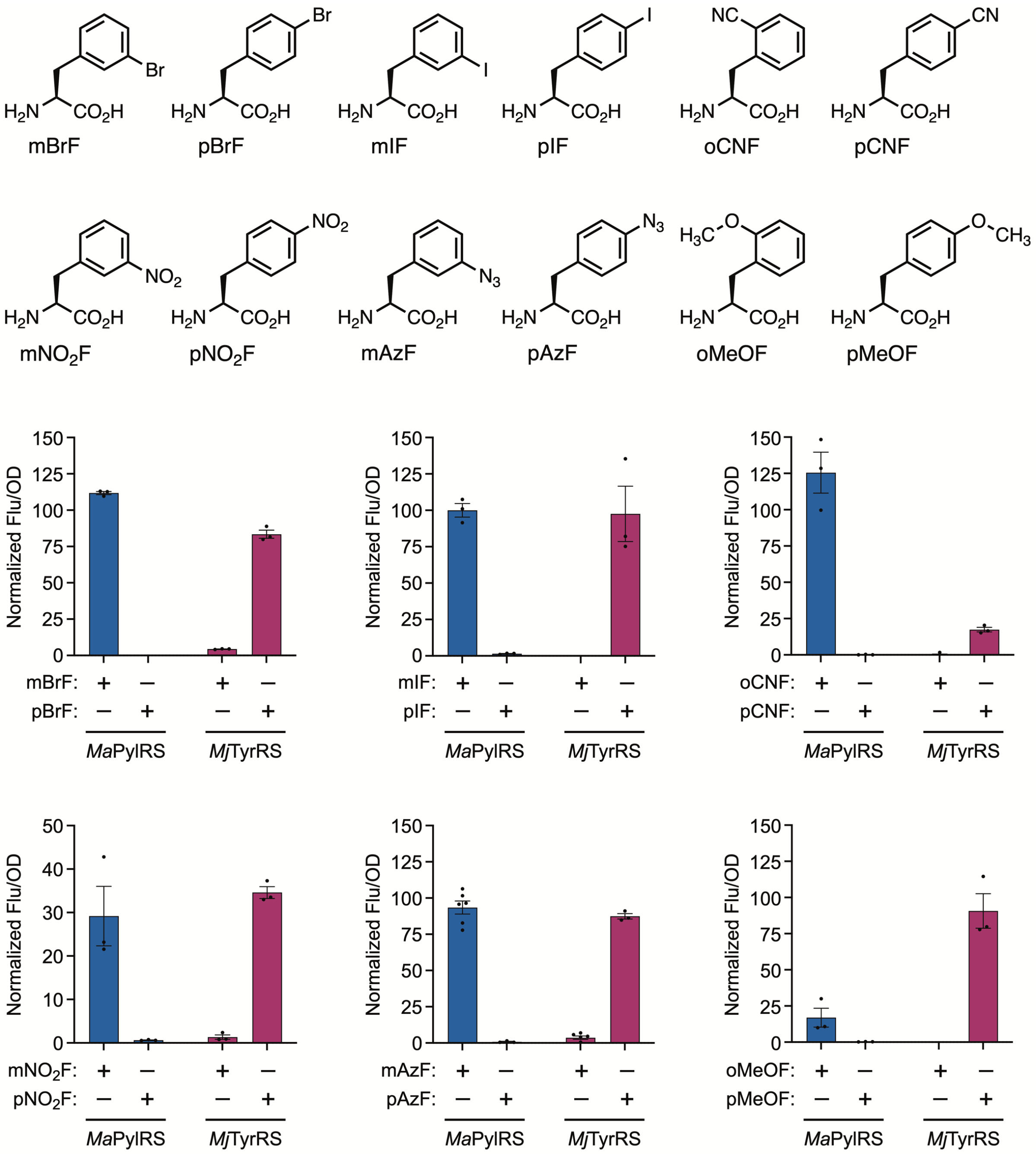


**Figure S14. Substrate specificities of *Ma*PylRS(N166S) and AzFRS.2.t1.** *Ma*PylRS(N166S) is selective for *ortho*- and *meta*-substituted phenylalanine derivatives while AzFRS.2.t1 is selective for *para*-substituted phenylalanine. (Top) Structures of amino acid with which we tested *Ma*PylRS(N166S) and AzFRs.2.t1 activity. (Bottom) Expression of sfGFP[2UAG] in DH10B cells co-expressing *Ma*PylRS(N166S) and *Ma*tRNA(6)$^{Pyl}$ or AzFRS.2.t1 and *i*tRNA$^{Ty2}$. Cells were grown in defined media supplemented with 0.1 mM IPTG, 0.2% arabinose, and 2 mM of the indicated amino acid. Data were collected 18 hours after induction and are displayed as the mean ± SEM of three biological replicates.

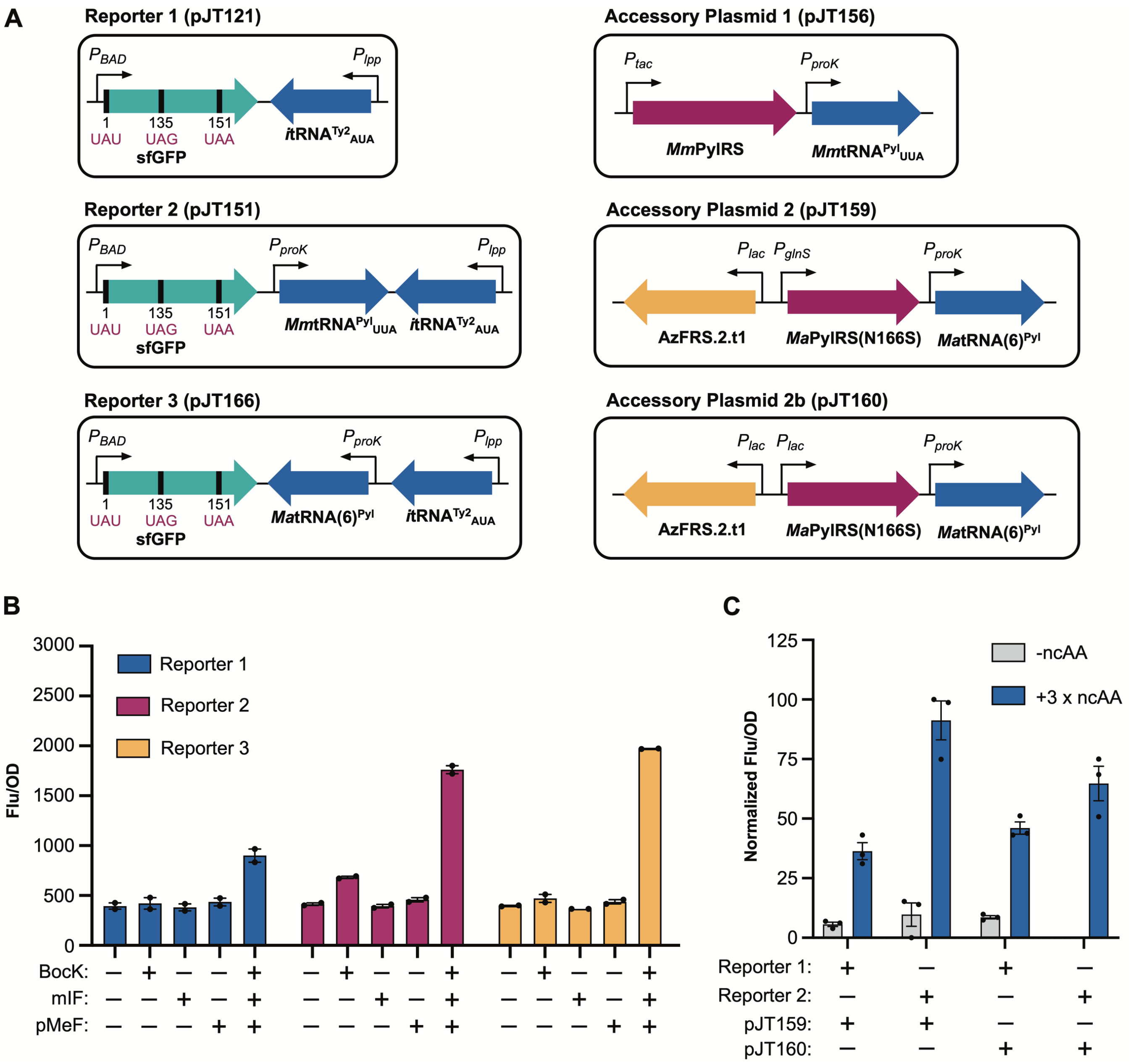


**Figure S15. Optimizing triple ncAA incorporation.** (A) Maps of the three-plasmid system used to simultaneously install three distinct ncAAs. Cells were co-transformed with one of three reporter plasmids (Reporters 1–3) and two accessory plasmids (pJT156 and either pJT159 or pJT160). Optimized expression was achieved in cells transformed with Reporter 3, pJT156, and pJT159. (B) Optimizing pyrrolysine tRNA expression. Adding the *Mm*tRNA$^{Pyl}_{UUA}$ or *Ma*tRNA(6)$^{Pyl}$ gene to the reporter plasmid significantly improved protein yield; however, when Reporter 2 was used, excess *Mm*tRNA$^{Pyl}_{UUA}$ suppressed both UAA and UAG (*see* Figure S16). Correct ncAA incorporation was achieved with Reporter 3. Data were collected 24 hours after induction and are displayed as the mean ± SEM of two biological replicates. (C) Optimizing *Ma*PylRS(N166S) expression. The reporter gene sfGFP[1UAU-135UAG-151UAA] was expressed from Reporter 1 or Reporter 2 in cells containing pJT156 and either pJT159 or pJT160. The latter is a variant of pJT159 in which *Ma*PylRS(N166S) was placed under a strong lac promoter. Data were collected 24 hours after induction and are displayed as the mean ± SEM of three biological replicates. In all cases, cells were grown in defined media[1] supplemented with 0.1 mM IPTG, 0.2% arabinose, and 2 mM each of BocK, mIF, and pMeF.

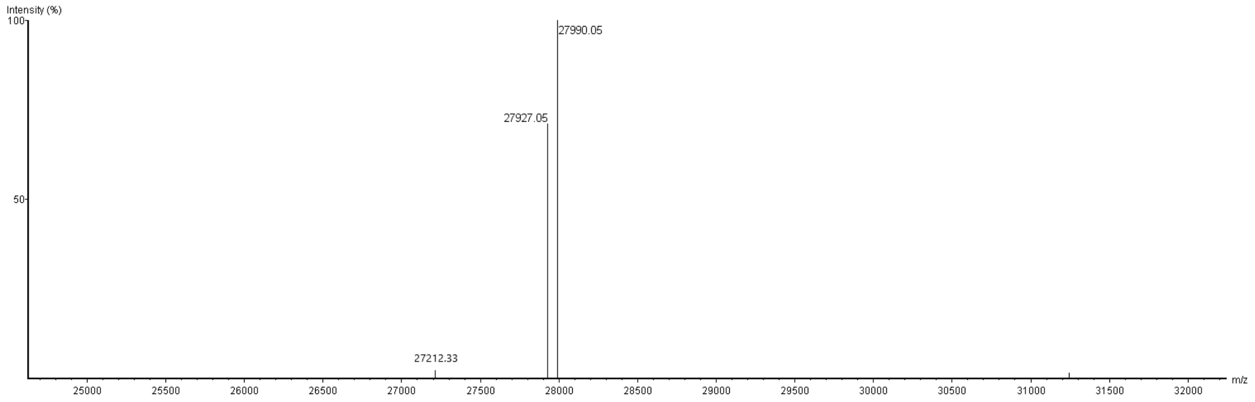


**Figure S16. LC-MS of sfGFP-1pAcF-135mIF-151PrK with misincorporation of PrK at position 135.** The protein was expressed from Reporter 2 in cells containing pJT156 and pJT159 (*see* Figure S15). The major peak corresponds to sfGFP with all three ncAAs at the desired positions (theoretical mass = 27992 Da; observed mass = 27990 Da). The minor peak corresponds to sfGFP-1pAcF-135PrK-151PrK (theoretical mass = 27929 Da; observed mass = 27927 Da). This product results from misincorporation of PrK in response to UAG, a consequence of excess *Mm*tRNA$^{Pyl}_{UUA}$. Misincorporation at UAG is not observed when *Ma*tRNA(6)$^{Pyl}$ is in excess.

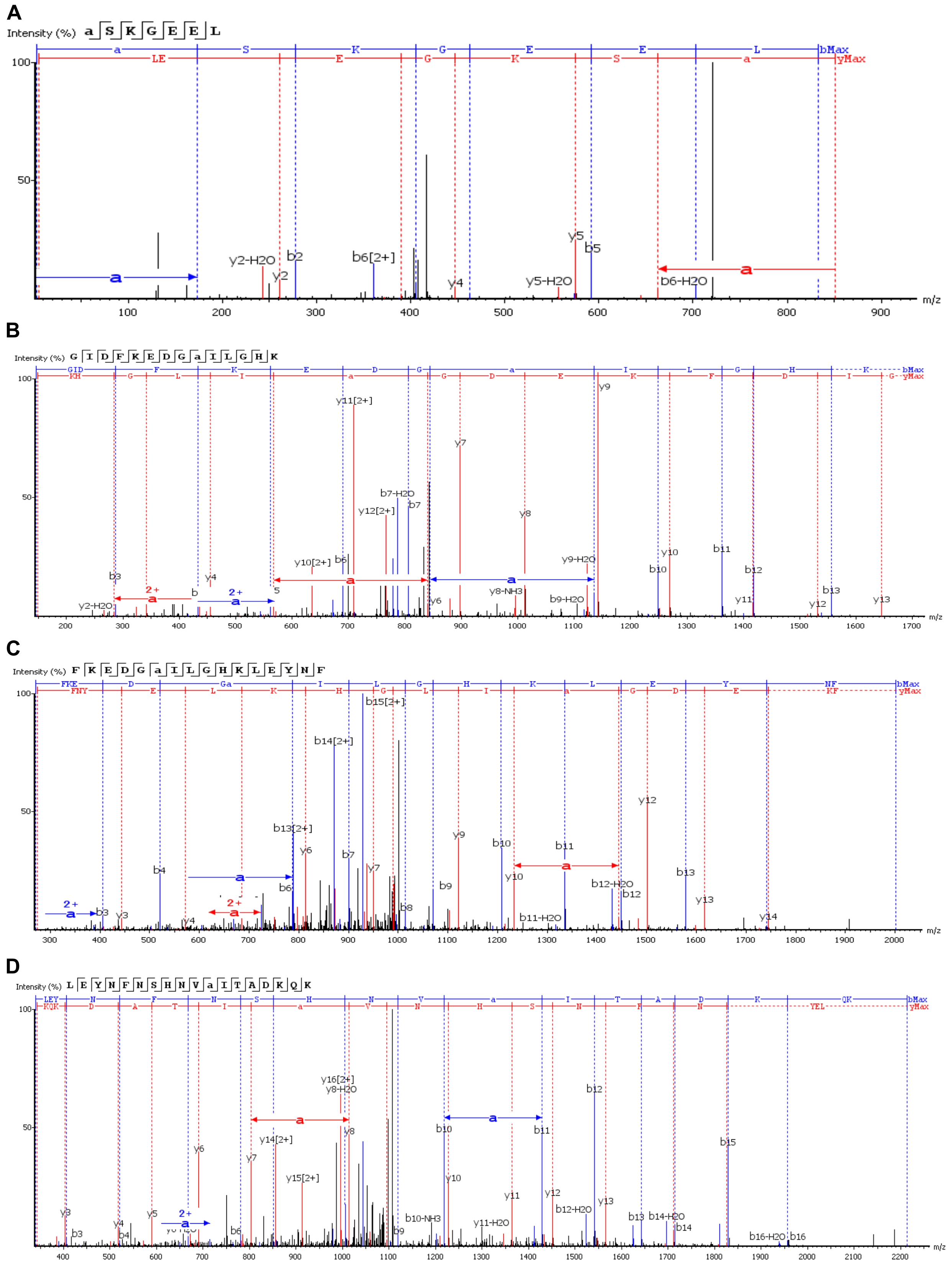

**Figure S17. MS/MS analysis of sfGFP-1pAcF-135mIF-151PrK with misincorporation of PrK at position 135.** The spectra support incorporation of pAcF at position 1 (A), mIF at position 135 (B), PrK at position 135 (C) and PrK at position 151 (D). For each spectrum, the position of the ncAA is indicated by the letter *a*.

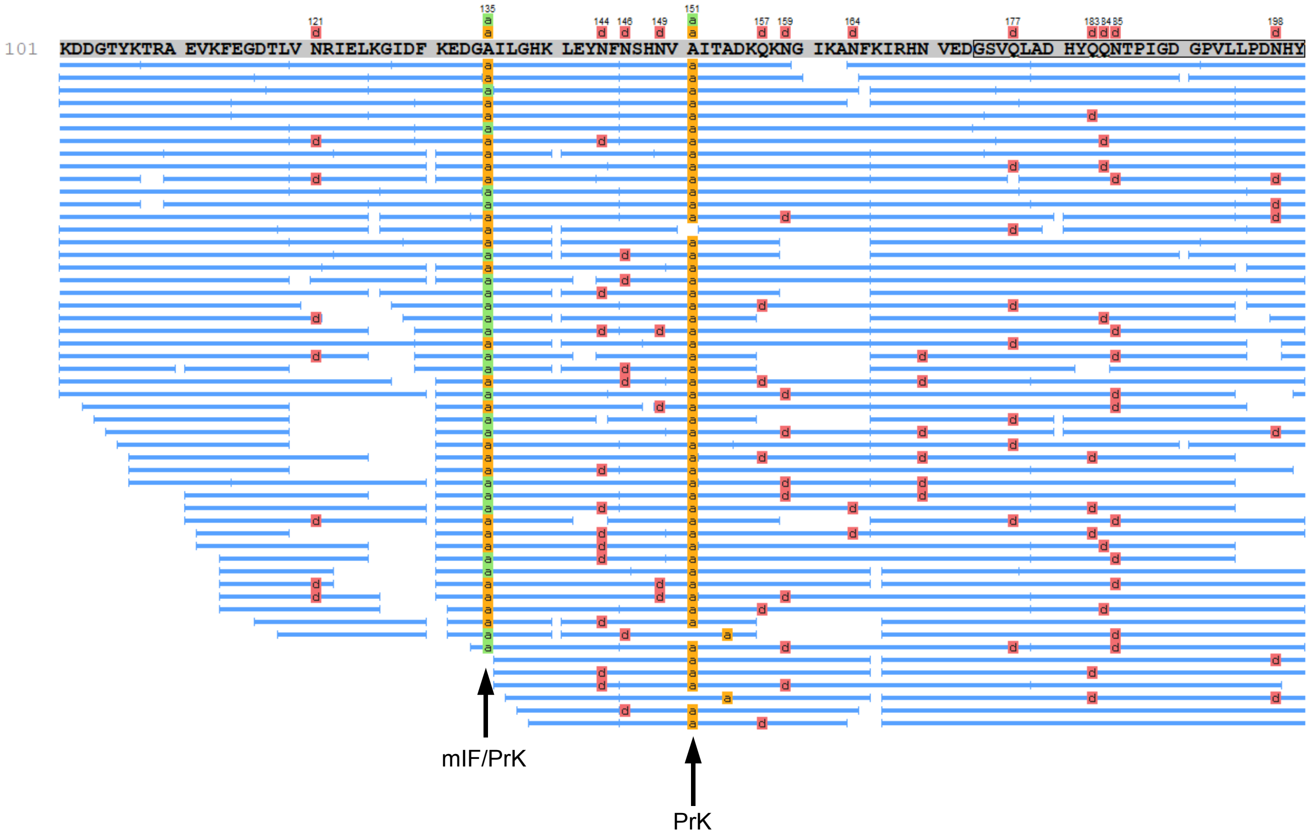


**Figure S18. Protein coverage map of sfGFP-1pAcF-135mIF-151PrK with misincorporation of PrK at position 135.** The map shows all peptides containing mIF and PrK that were identified during MS/MS analysis. A mixture of mIF and PrK is seen at position 135, whereas, only PrK was observed at position 151.

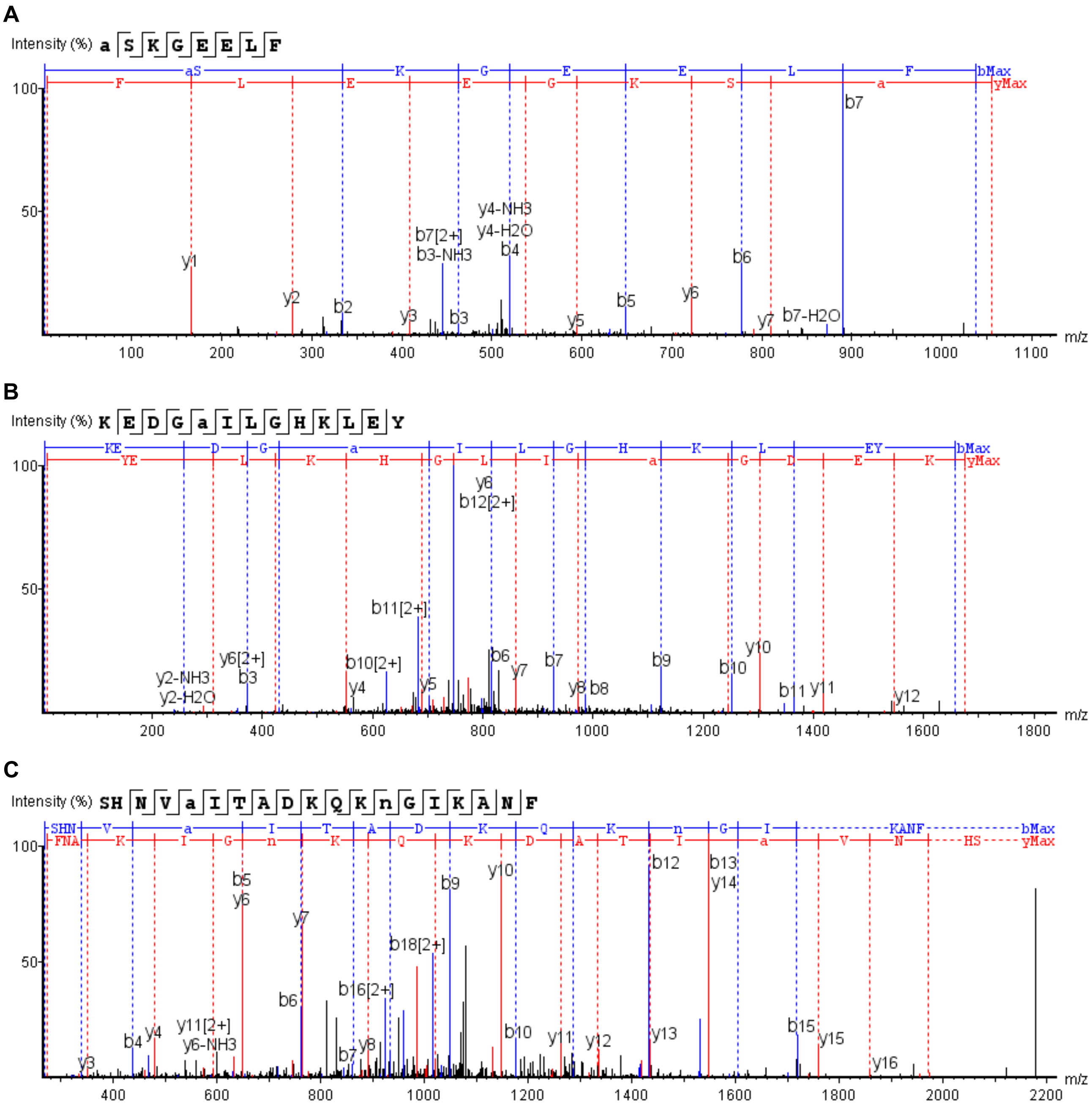


**Figure S19. MS/MS analysis of sfGFP-1pAcF-135mIF-151PrK.** The spectra support incorporation of pAcF at position 1 (A), mIF at position 135 (B), and PrK at position 151 (C). For each spectrum, the position of the ncAA is indicated by the letter *a*.

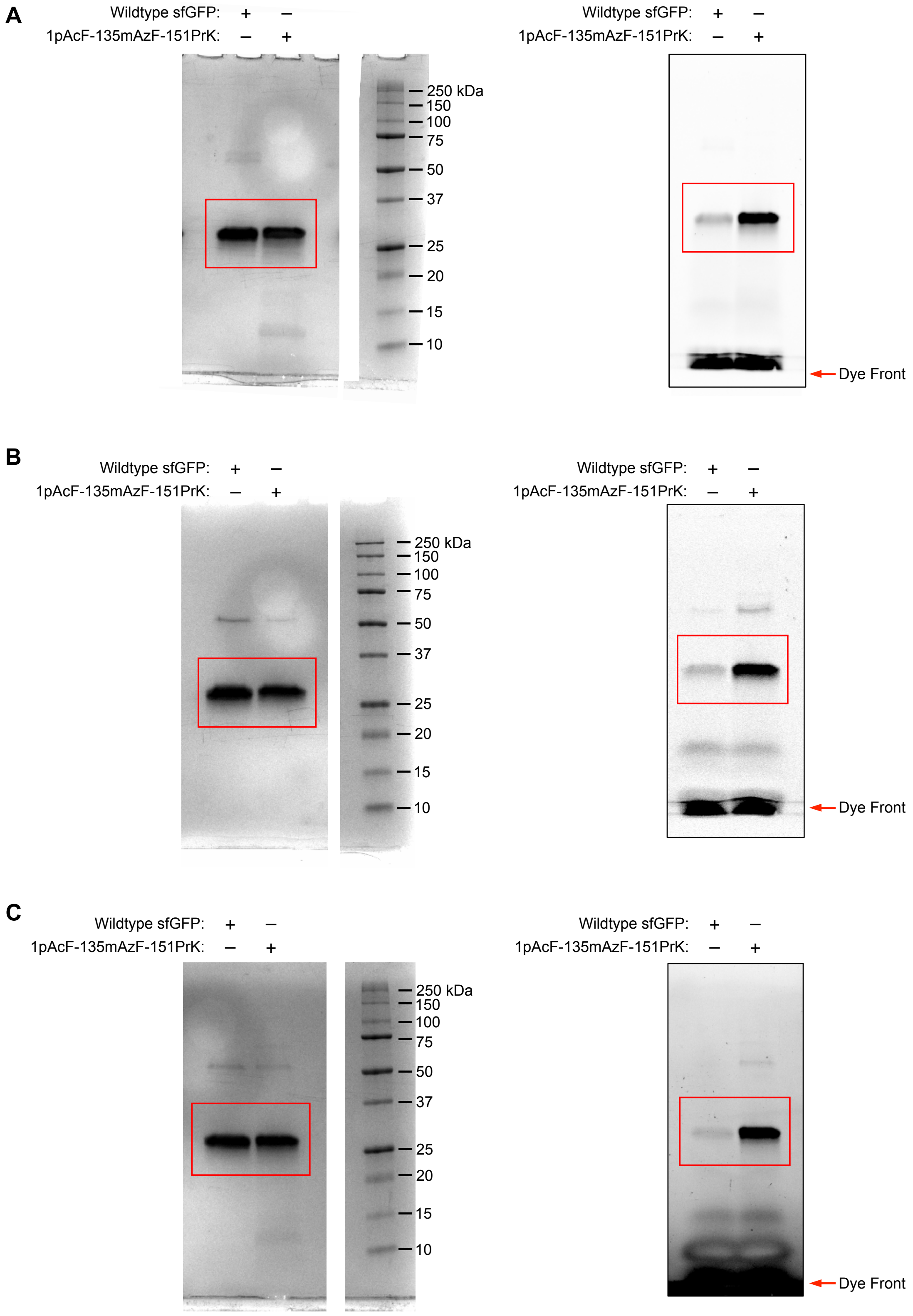


**Supplementary Figure 20. Extended data for Figure 5D.** Uncropped gel images for labeling of sfGFP-1pAcF-135mAzF-151PrK with Fluor 488-hydroxylamine (A), Fluor 488-alkyne (B), and coumarin azide (C). Coomassie stained gels are shown on the left, in-gel fluorescence images are shown on the right.

### Supplementary Tables

#### Table S1. Plasmids used in this study.

| Plasmid Name | Resistance | Origin | ORF 1 | | ORF2 | | ORF3 | | Figure(s) |
|---|---|---|---|---|---|---|---|---|---|
| | | | Prom | Gene | Prom | Gene | Prom | Gene | |
| **pCAM Backbone** | | | | | | | | | |
| pCAM-Ma | Tet | p15A | $P_{cat}$ | cat[112UAG] | $P_{Lpp}$ | *Ma*tRNA(6)$^{Pyl}_{CUA}$ | - | - | 4D, S11 |
| **pBAD Backbone** | | | | | | | | | |
| pJT008 | Amp | pBR322 | $P_{BAD}$ | sfGFP[2UAG] | $P_{Lpp}$ | *Mm*tRNA$^{Pyl}_{CUA}$ | – | – | 4B |
| pJT010 | Amp | pBR322 | $P_{BAD}$ | sfGFP[2UAG] | $P_{Lpp}$ | *Ma*tRNA(6)$^{Pyl}_{CUA}$ | – | – | 4C, S12-14 |
| pJT052 | Amp | pBR322 | $P_{BAD}$ | sfGFP[1UAG] | $P_{Lpp}$ | *i*tRNA$^{Ty2}_{CUA}$ | – | – | 2B, 2C |
| pJT055 | Amp | pBR322 | $P_{BAD}$ | sfGFP[2UAG] | $P_{Lpp}$ | *i*tRNA$^{Ty2}_{CUA}$ | – | – | 4A, S14 |
| pJT100-pJT107[a] | Amp | pBR322 | $P_{BAD}$ | sfGFP[1NNN] | $P_{Lpp}$ | *i*tRNA$^{Ty2}_{CUA}$ | – | – | 2B |
| pJT108-pJT115[b] | Amp | pBR322 | $P_{BAD}$ | sfGFP[1NNN] | $P_{Lpp}$ | *i*tRNA$^{Ty2}_{NNN}$ | – | – | 2C-E, S2A, S3-4 |
| pJT117 | Amp | pBR322 | $P_{BAD}$ | sfGFP$^{Opt}$[1UAU] | $P_{Lpp}$ | *i*tRNA$^{Ty2}_{AUA}$ | – | – | 2G, S1, S2B, S5 |
| pJT121 | Amp | pBR322 | $P_{BAD}$ | sfGFP$^{Opt}$[1UAU, 135UAG, 151UAA] | $P_{Lpp}$ | *i*RNA$^{Ty2}_{AUA}$ | – | – | S15A-C |
| pJT123 | Amp | pBR322 | $P_{BAD}$ | sfGFP$^{Opt}$[2UAG] | $P_{Lpp}$ | *i*RNA$^{Ty2}_{AUA}$ | – | – | S5 |
| pJT124 | Amp | pBR322 | $P_{BAD}$ | sfGFP$^{Opt}$[2UAA] | $P_{Lpp}$ | *i*RNA$^{Ty2}_{AUA}$ | – | – | S5 |
| pJT151 | Amp | pBR322 | $P_{BAD}$ | sfGFP$^{Opt}$[1UAU, 135UAG, 151UAA] | $P_{Lpp}$ | *i*RNA$^{Ty2}_{AUA}$ | $P_{ProK}$ | *Mm*tRNA$^{Pyl}_{UUA}$ | S15A-C, S16-18 |
| pJT166 | Amp | pBR322 | $P_{BAD}$ | sfGFP$^{Opt}$[1UAU, 135UAG, 151UAA] | $P_{Lpp}$ | *i*RNA$^{Ty2}_{AUA}$ | $P_{ProK}$ | *Ma*tRNA(6)$^{Pyl}_{CUA}$ | 5A-D, S15A-B, S19-20 |
| pJT171[c] | Amp | pBR322 | $P_{BAD}$ | sfGFP$^{Opt}$[1UAU, 151UAA] | $P_{Lpp}$ | *i*RNA$^{Ty2}_{AUA}$ | $P_{ProK}$ | *Mm*tRNA$^{Pyl}_{UUA}$ | 3A, 3C, 3E, S6A, S6C, S8-9 |
| pJT172[d] | Amp | pBR322 | $P_{BAD}$ | sfGFP$^{Opt}$[1UAU, 135UAG] | $P_{Lpp}$ | *i*RNA$^{Ty2}_{AUA}$ | $P_{ProK}$ | *Ma*tRNA(6)$^{Pyl}_{CUA}$ | 3B, 3D, 3E, S6B, S6D, S7, S9 |
| **pMW Backbone** | | | | | | | | | |
| pJT048 | Spec | SC101 | $P_{Lac}$ | pCNFRS | – | – | – | – | 2B-E, S2A, S3-4 |
| pJT057 | Spec | SC101 | $P_{Lac}$ | AzFRS.2.t1 | – | – | – | – | 2G, 4A, S1, S2B, S5, S14 |
| pJT064 | Spec | SC101 | $P_{Lac}$ | *Mm*PylRS | – | – | – | – | 4B |
| pJT065 | Spec | SC101 | $P_{Lac}$ | *Ma*PylRS | – | – | – | – | 4C |
| pJT131 | Spec | SC101 | $P_{Lac}$ | *Mm*PylRS(N346A, C348A) | – | – | – | – | 4B |
| pJT135 | Spec | SC101 | $P_{Lac}$ | *Ma*PylRS(N166V) | – | – | – | – | S12 |
| pJT136 | Spec | SC101 | $P_{Lac}$ | *Ma*PylRS(N166S, V168I) | – | – | – | – | S12 |
| pJT137 | Spec | SC101 | $P_{Lac}$ | *Ma*PylRS(N166G, V168I) | – | – | – | – | S12 |
| pJT138 | Spec | SC101 | $P_{Lac}$ | *Ma*PylRS(N166S) | – | – | – | – | 4C, S12-14 |

| | | | | | | | | | |
|---|---|---|---|---|---|---|---|---|---|
| **pULTRA Backbone** | | | | | | | | | |
| pJT156 | Spec | CloDF13 | $P_{tac}$ | *Mm*PylRS | $P_{proK}$ | *Mm*tRNA$^{Pyl}_{UUA}$ | – | – | 3A, 3C, 3E, 5A-D, S6A, S6C, S8-9, S15A-C, S16-20 |
| pJT173 | Spec | CloDF13 | $P_{tac}$ | *Ma*PylRS | $P_{proK}$ | *Ma*tRNA(6)$^{Pyl}_{CUA}$ | – | – | 3B, 3D-E, S6B, S6D, S7, S9 |
| **pSTART Backbone** | | | | | | | | | |
| pJT159 | Cm | SC101 | $P_{Lac}$ | AzFRS.2.t1 | $P_{glnS}$ | *Ma*PylRS(N166S) | $P_{ProK}$ | *Ma*tRNA(6)$^{Pyl}_{CUA}$ | 5A-D, S15A-C, S16-20 |
| pJT160 | Cm | SC101 | $P_{Lac}$ | AzFRS.2.t1 | $P_{Lac}$ | *Ma*PylRS(N166S) | $P_{ProK}$ | *Ma*tRNA(6)$^{Pyl}_{CUA}$ | S15A, S15C |
| pJT174 | Cm | SC101 | $P_{Lac}$ | AzFRS.2.t1 | – | – | – | – | 3A-E, S6A-D, S7-9 |

[a]sfGFP variants with initiating codons UAC, UAU, UUC, UGC, UCC, AAC, GAC, and CAC.
[b]sfGFP variants with initiating codons UAC, UAU, UUC, UGC, UCC, AAC, GAC, and CAC and corresponding *i*tRNA$^{Ty2}$ anticodon mutants.
[c]ORF4: Promoter, $P_{ProK}$; Gene, *Ma*tRNA(6)$^{Pyl}_{CUA}$
[d]ORF4: Promoter, $P_{ProK}$; Gene, *Mm*tRNA$^{Pyl}_{UUA}$

#### Table S2. Oligonucleotides used in this study

| Name | Sequence (5'→3') |
|---|---|
| **JT27** | GCGAAGGCGAAGCGGAAGCTTAAAAAAAATCCTTAGCTTT |
| **JT28** | GACAGGCACATTATGCTGGCGCCGCTTCTTTGA |
| **JT169** | GCGGCCGCACCTCCTTTG |
| **JT176** | TAAAACCTAGCATAGCGGGGTTCGACACCCCGGTCTCTCGCCAAATTCGAAAAGCCTGCTCAACG |
| **JT261** | TTTAGCTTCCTCCTGTTAGCCCA |
| **JT262** | CAGGAGGAAGCTAAATATGACAAGGGCGAAGAACTG |
| **JT263** | CAGGAGGAAGCTAAATTCGACAAGGGCGAAGAACTG |
| **JT264** | CAGGAGGAAGCTAAATGCGACAAGGGCGAAGAACTG |
| **JT265** | CAGGAGGAAGCTAAATCCGACAAGGGCGAAGAACTG |
| **JT266** | CAGGAGGAAGCTAAAAACGACAAGGGCGAAGAACTG |
| **JT267** | CAGGAGGAAGCTAAAGACGACAAGGGCGAAGAACTG |
| **JT268** | CAGGAGGAAGCTAAACACGACAAGGGCGAAGAACTG |
| **JT269** | CAGCATGGTAAATTCTTCCAGATGCATACCG |
| **JT270** | GAATTTACCATGCTGNNMCTGNNMGATATGGGTCCGCGTGGTG |
| **JT271** | AACCCGAAGATCGTCGGTTCAAATCCG |
| **JT272** | GACGATCTTCGGGTTTACAGCCCGACGAGCTACCA |
| **JT273** | GACGATCTTCGGGTTTATAGCCCGACGAGCTACCA |
| **JT274** | GACGATCTTCGGGTTTTCAGCCCGACGAGCTACCA |
| **JT275** | GACGATCTTCGGGTTTGCAGCCCGACGAGCTACCA |
| **JT276** | GACGATCTTCGGGTTTCCAGCCCGACGAGCTACCA |
| **JT277** | GACGATCTTCGGGTTAACAGCCCGACGAGCTACCA |
| **JT278** | GACGATCTTCGGGTTGACAGCCCGACGAGCTACCA |
| **JT279** | GACGATCTTCGGGTTCACAGCCCGACGAGCTACCA |
| **JT290** | CTCGAGATCTGCAGCTGGTACC |
| **JT291** | TTTAGCTTCCTCCTGTTAGCCC |
| **JT292** | CAGGAGGAAGCTAAATATAGCAAGGGCGAAGAGTTATTCA |
| **JT293** | GCTGCAGATCTCGAGTCAATGATGATGATGATGATGTGAG |
| **JT294** | AGACGGATAGATTTTAGGTCATAAGCTGGAGTAC |
| **JT295** | AAAATCTATCCGTCTTCTTTGAAATCAATGCC |
| **JT296** | CAACGTATAAATTACAGCTGACAAACAGAAGAACG |
| **JT297** | GTAATTTATACGTTGTGAGAGTTAAAATTGTAC |
| **JT298** | CTAAAATGTAAAAGGGCGAAGAGTTATTCACAGGG |
| **JT299** | CCTTTTACATTTTAGCTTCCTCCTGTTAGCCC |
| **JT300** | CTAAAATGTAGAAGGGCGAAGAGTTATTCACAGGG |
| **JT301** | CCTTCTACATTTTAGCTTCCTCCTGTTAGCCC |
| **JT302** | CAGGAGGAAGCTAAATAGAGCAAGGGCGAAGAGTTATTCA |
| **JT313** | GCAGCAGATCAATTCGCGC |
| **JT314** | CTCGCGCGTTTCGGTGATG |
| **JT317** | ACCGAAACGCGCGAGAGGCATTTTGCTATTAAGGGATTG |
| **JT318** | GAATTGATCTGCTGCGCATGCAAAAAAGCCTGC |
| **JT347** | AGATTTTAGAGCCAATTAGAAAGAG |
| **JT348** | AGGAGGTGCGGCCGCATGGATAAAAAACCGCTGAATACCC |
| **JT349** | TTGGCTCTAAAATCTTCACAGGTTGGTACTAATACCATTGTAA |
| **JT351** | CTGCAGTTTCAAACGCTAAATTGCC |

| | |
|---|---|
| **JT352** | ATGGGATTCCTCAAAGCGTAAAC |
| **JT353** | TTTGAGGAATCCCATATGACAGTCAAATACACCGATGCAC |
| **JT354** | CGTTTGAAACTGCAGTCAATTGATCTTGGCACCATTCAG |
| **JT355** | CTATGCTAGGTTTTAGAGACCCGCTGGTCGCCGGACCGTCCCCCAATGCGGGGCGCATCTTACTG |
| **JT356** | CTTTAAATCCGTTCAGCCGGGTTAGATTCCCGGGGTTTCCGCCAAATTCGAAAAGCCTGCTCAACG |
| **JT357** | GCTGAACGGATTTAAAGTCCGTTCGATCTACATGATCAGGTTTCCAATGCGGGGCGCATCTTACTG |
| **JT360** | AGACGGAAACATTTTAGGTCATAAGCTGGAGTAC |
| **JT361** | AAAATGTTTCCGTCTTCTTTGAAATCAATGCC |
| **JT362** | CAACGTATACATTACAGCTGACAAACAGAAGAACG |
| **JT363** | GTAATGTATACGTTGTGAGAGTTAAAATTGTAC |
| **JT370** | CTCATCCTGTCTCTTGATCACTACC |
| **JT371** | CGAACTGAGATACCTACAGCG |
| **JT372** | AGGTATCTCAGTTCGGCTCAGTCGAAAGACTGGGCC |
| **JT373** | AAGAGACAGGATGAGGCGAAAATGAGACGTTGATCGGC |
| **JT386** | CTGCAGTTTCAAACGCTAAATTG |
| **JT387** | GGACAGGCTGACAACTGTTAG |
| **JT388** | GTTGTCAGCCTGTCCGCAGTGAGCGCAACGCAA |
| **JT389** | CGTTTGAAACTGCAGTCAATTGATCTTGGCACCATTCA |
| **JT394** | CGGCATCCGCTTACAGACAAG |
| **JT395** | GGAGCAGACAAGCCCGTC |
| **JT396** | GGGCTTGTCTGCTCCGCATGCAAAAAAGCCTGC |
| **JT397** | TGTAAGCGGATGCCGAGGCATTTTGCTATTAAGGGATTG |
| **JT398** | TTTAATATGAGCAGCTCAGGGTCGAATTTG |
| **JT399** | GCTGCTCATATTAAAGCGGGACAGGCTGAC |
| **407.F** | GAATCCTTCCCCCACCACCAGGCATAAGCTTGGCGTAATCA |
| **407.R** | GGCCGCTCGGGAACCCCACCTATAGTGAGTCGTATTAGGATCCCCGGGTACC |
| **413.F** | GGTGGGGTTCCCGAGCGGCCAAAGGGAGCAGACTGTAAATCTGCCGTCATCGACTTCGAAGGTTCGAATCCTTCCCCCACCACCA |
| **413.R** | TGGTGGTGGGGGAAGGATTCGAACCTTCGAAGTCGATGACGGCAGATTTACAGTCTGCTCCCTTTGGCCGCTCGGGAACCCCACC |
| **417.F** | TCCTAATACGACTCACTATACGCGGGGTGGAGCAGCCTGGTAGCTCGTCGGGCTATAAACCCGAAGATCGTCGGTTCAAATCCGGC CCCCGCGACCATGCATAAGCTTGGCCTGGCT |
| **417.R** | AGCCAGGCCAAGCTTATGCATGGTCGCGGGGGCCGGATTTGAACCGACGATCTTCGGGTTTATAGCCCGACGAGCTACCAGGCTG CTCCACCCCGCGTATAGTGAGTCGTATTAGGA |
| **418.F** | AAATCCGGCCCCCGCGACCATGCATAAGCTTGGCC |
| **418.R** | CCAGGCTGCTCCACCCCGCGTATAGTGAGTCGTATTAGGATCCCCGG |
| **436** | TGGTCGCGGGGGCCGGATTTG |
| **437** | TGGTGGTGGGGGAAGG |
| **pUC18 _un** | CGACGTTGTAAAACGACGGC |

**DNA Sequences**

**a.** sfGFP[1UAU]: The initiating codon is highlighted; internal UAU codons are shown in bold.

TATGACAAGGGCGAAGAACTGTTTACGGGCGTGGTGCCGATTCTGGTGGAACTGGATGGTGATGT
CAATGGTCACAAATTCAGCGTGCGCGGCGAAGGTGAAGGCGATGCAACCAATGGTAAACTGACGC
TGAAGTTTATTTGCACCACGGGTAAACTGCCGGTTCCGTGGCCGACCCTGGTCACCACGCTGACG
**TAT**GGTGTTCAGTGTTTCAGTCGTTACCCGGATCACATGAAACGCCACGACTTTTTCAAGTCCGCG
ATGCCGGAAGGT**TAT**GTCCAAGAACGTACCATCTCATTTAAAGATGACGGCACCTACAAAACGCGC
GCCGAAGTGAAATTCGAAGGTGATACGCTGGTTAACCGTATTGAACTGAAAGGCATCGATTTTAAG
GAAGACGGTAATATTCTGGGCCATAAACTGGAA**TAT**AACTTCAATTCGCACAACGTGTACATCACC
GCAGATAAGCAGAAGAACGGTATCAAGGCTAACTTCAAGATCCGCCATAATGTGGAAGATGGCAG
CGTTCAACTGGCCGACCAC**TAT**CAGCAAAACACCCCGATTGGTGATGGCCCGGTCCTGCTGCCG
GACAATCATTACCTGAGCACGCAGTCTGTGCTGAGTAAAGATCCGAACGAAAAGCGTGACCACAT
GGTCCTGCTGGAATTCGTGACCGCGGCCGGCATCACGCACGGTATGGACGAACTG**TAT**AAAGGC
TCACATCATCATCATCATCATTGA

**b.** $sfGFP^{Opt}$[1UAU]: The initiating codon is highlighted.

TATAGCAAGGGCGAAGAGTTATTCACAGGGGTTGTCCCTATTCTTGTAGAATTGGATGGAGACGTA
AACGGCCATAAATTCTCCGTACGCGGGGAAGGCGAAGGTGATGCTACAAATGGGAAATTAACTCT
GAAATTTATTTGCACAACCGGAAAATTACCTGTGCCTTGGCCTACTCTGGTCACGACCTTAACATAC
GGTGTACAGTGCTTTTCACGTTACCCAGATCATATGAAGCGCCACGATTTCTTCAAGAGTGCAATG
CCTGAGGGGTACGTTCAAGAACGTACCATTTCGTTTAAGGACGATGGAACCTACAAAACCCGTGC
AGAGGTCAAGTTTGAAGGTGACACTTTAGTTAACCGTATTGAATTAAAGGGCATTGATTTCAAAGAA
GACGGAAACATTTTAGGTCATAAGCTGGAGTACAATTTTAACTCTCACAACGTATACATTACAGCTG
ACAAACAGAAGAACGGTATTAAAGCGAATTTCAAGATCCGTCATAACGTAGAAGATGGAAGCGTAC
AATTGGCGGATCACTACCAACAGAATACTCCTATCGGCGACGGCCCTGTGCTGCTTCCCGACAAC
CATTACTTGTCCACTCAGAGCGTATTATCAAAGGACCCAAACGAGAAACGCGACCATATGGTGCTT
TTAGAGTTCGTGACCGCCGCTGGCATCACCCATGGTATGGACGAGCTGTACAAGGGCTCACATCA
TCATCATCATCATTGA

**c.** $sfGFP^{Opt}$[1UAU-135UAG-151UAA]: The initiating codon is highlighted; internal nonsense codons are shown in bold.

TATAGCAAGGGCGAAGAGTTATTCACAGGGGTTGTCCCTATTCTTGTAGAATTGGATGGAGACGTA
AACGGCCATAAATTCTCCGTACGCGGGGAAGGCGAAGGTGATGCTACAAATGGGAAATTAACTCT
GAAATTTATTTGCACAACCGGAAAATTACCTGTGCCTTGGCCTACTCTGGTCACGACCTTAACATAC
GGTGTACAGTGCTTTTCACGTTACCCAGATCATATGAAGCGCCACGATTTCTTCAAGAGTGCAATG
CCTGAGGGGTACGTTCAAGAACGTACCATTTCGTTTAAGGACGATGGAACCTACAAAACCCGTGC
AGAGGTCAAGTTTGAAGGTGACACTTTAGTTAACCGTATTGAATTAAAGGGCATTGATTTCAAAGAA
GACGGA**TAG**ATTTTAGGTCATAAGCTGGAGTACAATTTTAACTCTCACAACGTA**TAA**ATTACAGCTG
ACAAACAGAAGAACGGTATTAAAGCGAATTTCAAGATCCGTCATAACGTAGAAGATGGAAGCGTAC
AATTGGCGGATCACTACCAACAGAATACTCCTATCGGCGACGGCCCTGTGCTGCTTCCCGACAAC
CATTACTTGTCCACTCAGAGCGTATTATCAAAGGACCCAAACGAGAAACGCGACCATATGGTGCTT
TTAGAGTTCGTGACCGCCGCTGGCATCACCCATGGTATGGACGAGCTGTACAAGGGCTCACATCA
TCATCATCATCATTGA

**d.** *i*$tRNA^{Ty2}_{AUA}$:

CGCGGGGTGGAGCAGCCTGGTAGCTCGTCGGGCTATAAACCCGAAGATCGTCGGTTCAAATCCG
GCCCCCGCGACCA

**e.** *Mm*$tRNA^{Pyl}_{UUA}$:

GGAAACCTGATCATGTAGATCGAACGGACTTTAAATCCGTTCAGCCGGGTTAGATTCCCGGGGTTT
CCGCCA

**f.** *Ma*$tRNA(6)^{Pyl}_{CUA}$:

GGGGGACGGTCCGGCGACCAGCGGGTCTCTAAAACCTAGCATAGCGGGGTTCGACACCCCGGTC
TCTCGCCA

**g**. *Mj*TyrRS variant pCNFRS:

ATGGACGAATTTGAAATGATAAAGAGAAACACATCTGAAATTATCAGCGAGGAAGAGTTAAGAGAG
GTTTTAAAAAAAGATGAAAAATCTGCTCTGATAGGTTTTGAACCAAGTGGTAAAATACATTTAGGGC
ATTATCTCCAAATAAAAAAGATGATTGATTTACAAAATGCTGGATTTGATATAATTATAGTTTTGGCT
GATTTACATGCCTATTTAAACCAGAAAGGAGAGTTGGATGAGATTAGAAAAATAGGAGATTATAACA
AAAAAGTTTTTGAAGCAATGGGGTTAAAGGCAAAATATGTTTATGGAAGTGAATGGATGCTTGATAA
GGATTATACACTGAATGTCTATAGATTGGCTTTAAAAACTACCTTAAAAAGAGCAAGAAGGAGTATG
GAACTTATAGCAAGAGAGGATGAAAATCCAAAGGTTGCTGAAGTTATCTATCCAATAATGCAGGTT
AATGGTGCTCATTATCTTGGCGTTGATGTTGCAGTTGGGGGGATGGAGCAGAGAAAAATACACAT
GTTAGCAAGGGAGCTTTTACCAAAAAAGGTTGTTTGTATTCACAACCCTGTCTTAACGGGTTTGGAT
GGAGAAGGAAAGATGAGTTCTTCAAAAGGGAATTTTATAGCTGTTGATGACTCTCCAGAAGAGATT
AGGGCTAAGATAAAGAAAGCATACTGCCCAGCTGGAGTTGTTGAAGGAAATCCAATAATGGAGATA
GCTAAATACTTCCTTGAATATCCTTTAACCATAAAAAGGCCAGAAAAATTTGGTGGAGATTTGACAG
TTAATAGCTATGAGGAGTTAGAGAGTTTATTTAAAAATAAGGAATTGCATCCAATGGATTTAAAAAAT
GCTGTAGCTGAAGAACTTATAAAGATTTTAGAGCCAATTAGAAAGAGATTATAA

**h.** *Mj*TyrRS variant AzFRS.2.t1:

ATGGACGAATTTGAAATGATAAAGAGAAACACATCTGAAATTATCAGCGAGGAAGAGTTAAGAGAG
GTTTTAAAAAAGGATGAAAAATCTGCTCTGATAGGTTTTGAACCAAGTGGTAAAATACATTTAGGGC
ATTATCTCCAAATAAAAAAGATGATTGATTTACAAAATGCTGGATTTGATATAATTATATTGTTGGCT
GATTTACACGCCTATTTAAACCAGAAAGGAGAGTTGGATGAGATTAGAAAAATAGGAGATTATAACA
AAAAAGTTTTTGAAGCAATGGGGTTAAAGGCAAAATATGTTTATGGAAGTACTTATATGCTTGATAA
GGATTATACACTGAATGTCTATAGATTGGCTTTAAAAACTACCTTAAAAAGGGCAAGAAGGAGTATG
GAACTTATAGCAAGAGAGGATGAAAATCCAAAGGTTGCTGAAGTTATCTATCCAATAATGCAGGTT
AATGGTTGTCATTATAGGGGCGTTGATGTTGCTGTTGGAGGGATGGAGCAGAGAAAAATACACAT
GTTAGCAAGGGAGCTTTTACCAAAAAAGGTTGTTTGTATTCACAACCCTGTCTTAACGGGTTTGGAT
GGAGAAGGAAAGATGAGTTCTTCAAAAGGGAATTTTATAGCTGTTGATGACTCTCCAGAAGAGATT
AGGGCTAAGATAAAGAAAGCATACTGCCCAGCTGGAGTTGTTGAAGGAAATCCAATAATGGAGATA
GCTAAATACTTCCTTGAATATCCTTTAACCATAAAAGGGCCAGAAAAATTTGGTGGAGATTTGACAG
TTAATAGCTATGAGGAGTTAGAGAGTTTATTTAAAAATAAGGAATTGCATCCAATGCGCTTAAAAAA
TGCTGTAGCTGAAGAACTTATAAAGATTTTAGAGCCAATTAGAAAGAGATTATAA

**i.** *Ma*PylRS(N166S):

ATGACAGTCAAATACACCGATGCACAGATTCAGCGTCTGCGTGAATATGGTAATGGCACCTATGAA
CAGAAAGTGTTTGAAGATCTGGCAAGCCGTGATGCAGCATTTAGCAAAGAAATGAGCGTTGCAAG
CACCGACAATGAGAAAAAAATCAAAGGCATGATTGCAAACCCGAGCCGTCATGGTCTGACCCAGC
TGATGAATGATATTGCAGATGCACTGGTTGCCGAAGGTTTTATTGAAGTTCGTACCCCGATTTTCAT
CAGCAAAGATGCCCTGGCACGTATGACCATTACCGAAGATAAACCGCTGTTCAAACAGGTGTTTTG
GATTGATGAAAAACGTGCACTGCGTCCGATGCTGGCACCGAATCTGTATAGCGTTATGCGTGATCT
GCGCGATCATACCGATGGTCCGGTTAAAATCTTTGAAATGGGTAGCTGCTTTCGCAAAGAAAGCCA
TAGCGGTATGCATCTGGAAGAATTTACCATGCTGTCACTGGTAGATATGGGTCCGCGTGGTGATG

CAACCGAAGTTCTGAAAAACTATATTAGCGTTGTGATGAAAGCAGCAGGTCTGCCGGATTATGATC
TGGTTCAAGAAGAAAGCGACGTCTACAAAGAAACCATTGATGTGGAAATTAACGGCCAAGAAGTTT
GTAGCGCAGCAGTTGGTCCGCATTATCTGGATGCAGCACATGATGTGCATGAACCGTGGTCAGGT
GCAGGTTTTGGTCTGGAACGTCTGCTGACCATTCGTGAGAAATATAGCACCGTTAAAAAAGGTGGT
GCGAGCATTAGCTATCTGAATGGTGCCAAGATCAATTGA

**j.** *Mm*PylRS:

ATGGATAAAAAACCACTAAACACTCTGATATCTGCAACCGGGCTCTGGATGTCCAGGACCGGAACA
ATTCATAAAATAAAACACCACGAAGTCTCTCGAAGCAAAATCTATATTGAAATGGCATGCGGTGACC
ACCTTGTTGTAAACAACTCCAGGAGCAGCAGGACTGCAAGAGCGCTCAGGCACCACAAATACAGG
AAGACCTGCAAACGCTGCAGGGTTTCGGATGAGGATCTCAATAAGTTCCTCACAAAGGCAAACGA
AGACCAGACAAGCGTAAAAGTCAAGGTCGTTTCTGCCCCTACCAGAACGAAAAAGGCAATGCCAA
AATCCGTTGCGAGAGCCCCGAAACCTCTTGAGAATACAGAAGCGGCACAGGCTCAACCTTCTGGA
TCTAAATTTTCACCTGCGATACCGGTTTCCACCCAAGAGTCAGTTTCTGTCCCGGCATCTGTTTCAA
CATCAATATCAAGCATTTCTACAGGAGCAACTGCATCCGCACTGGTAAAAGGGAATACGAACCCCA
TTACATCCATGTCTGCCCCTGTTCAGGCAAGTGCCCCCGCACTTACGAAGAGCCAGACTGACAGG
CTTGAAGTCCTGTTAAACCCAAAAGATGAGATTTCCCTGAATTCCGGCAAGCCTTTCAGGGAGCTT
GAGTCCGAATTGCTCTCTCGCAGAAAAAAAGACCTGCAGCAGATCTACGCGGAAGAAAGGGAGAA
TTATCTGGGGAAACTCGAGCGTGAAATTACCAGGTTCTTTGTGGACAGGGGTTTTCTGGAAATAAA
ATCCCCGATCCTGATCCCTCTTGAGTATATCGAAAGGATGGGCATTGATAATGATACCGAACTTTC
AAAACAGATCTTCAGGGTTGACAAGAACTTCTGCCTGAGACCCATGCTTGCTCCAAACCTTTACAA
CTACCTGCGCAAGCTTGACAGGGCCCTGCCTGATCCAATAAAAATTTTTGAAATAGGCCCATGCTA
CAGAAAAGAGTCCGACGGCAAAGAACACCTCGAAGAGTTTACCATGCTGGCGTTCGCGCAGATGG
GATCGGGATGCACACGGGAAAATCTTGAAAGCATAATTACGGACTTCCTGAACCACCTGGGAATT
GATTTCAAGATCGTAGGCGATTCCTGCATGGTCTATGGGGATACCCTTGATGTAATGCACGGAGAC
CTGGAACTTTCCTCTGCAGTAGTCGGACCCATACCGCTTGACCGGGAATGGGGTATTGATAAACC
CTGGATAGGGGCAGGTTTCGGGCTCGAACGCCTTCTAAAGGTTAAACACGACTTTAAAAATATCAA
GAGAGCTGCAAGGTCCGAGTCTTACTATAACGGGATTTCTACCAACCTGTAA
